# Mining zebrafish microbiota reveals key community-level resistance against fish pathogen infection

**DOI:** 10.1101/2020.04.23.058222

**Authors:** Franziska A. Stressmann, Joaquin Bernal-Bayard, David Perez-Pascual, Bianca Audrain, Olaya Rendueles, Valérie Briolat, Sebastian Bruchmann, Stevenn Volant, Amine Ghozlane, Susanne Haussler, Eric Duchaud, Jean-Pierre Levraud, Jean-Marc Ghigo

**Affiliations:** Genetics of Biofilms Laboratory, Institut Pasteur, UMR CNRS2001, Paris 75015, France; Leibniz-Institute of Freshwater Ecology and Inland Fisheries, Department of Chemical Analytics and Biogeochemistry, Müggelseedamm 310, 12587, Berlin, Germany; Departamento de Genética, Facultad de Biología, Universidad de Sevilla, Apartado 1095, 41080 Sevilla, Spain; Microbial Evolutionary Genomics Laboratory, Institut Pasteur, UMR3525, Paris 75015, France; Macrophages and Development of Immunity Laboratory, Institut Pasteur, UMR3738 CNRS, Paris 75015, France; Department of Molecular Bacteriology, Helmholtz Centre for Infection Research, Braunschweig, Germany; University of Cambridge, Department of Veterinary Medicine, Madingley Road Cambridge CB3 0ES, UK; Hub de Bioinformatique et Biostatistique – Département Biologie Computationnelle, Institut Pasteur, USR 3756 CNRS, Paris, France; Department of Clinical Microbiology, Rigshospitalet; Copenhagen, 2100, Denmark; Unité VIM, INRAE, Université Paris-Saclay, 78350 Jouy-en-Josas, France

**Keywords:** Zebrafish, germ-free, probiotic, microbiota, colonization resistance, bacterial infection

## Abstract

The long-known resistance to pathogens provided by host-associated microbiota fostered the notion that adding protective bacteria could prevent or attenuate infection. However, the identification of endogenous or exogenous bacteria conferring such protection is often hindered by the complexity of host microbial communities. Here, we used zebrafish and the fish pathogen *Flavobacterium columnare* as a model system to study the determinants of microbiota-associated colonization resistance. We compared infection susceptibility in germ-free, conventional and re-conventionalized larvae and showed that a consortium of 10 culturable bacterial species are sufficient to protect zebrafish. Whereas survival to *F. columnare* infection does not rely on host innate immunity, we used antibiotic dysbiosis to alter zebrafish microbiota composition, leading to the identification of two different protection strategies. We first identified that the bacterium *Chryseobacterium massiliae* individually protects both larvae and adult zebrafish. We also showed that an assembly of 9 endogenous zebrafish species that do not otherwise protect individually confer a community-level resistance to infection. Our study therefore provides a rational approach to identify key endogenous protecting bacteria and promising candidates to engineer resilient microbial communities. It also shows how direct experimental analysis of colonization resistance in low-complexity *in vivo* models can reveal unsuspected ecological strategies at play in microbiota-based protection against pathogens.

## INTRODUCTION

Animal resident microbial consortia form complex and long-term associations with important community-level functions essential for host development and physiology (1, 2). Microbial ecosystems also provide protection against exogenous pathogens by inhibition of pathogen settlement and growth and/or stimulation of the host immune system (3–8). From the perspective of microbial community composition, a shift or reduction in resident microbial diversity, a phenomenon generally referred to as dysbiosis, is often associated with increased susceptibility to infection due to the loss or change in abundance of key microbial community members (3, 9). These observations early supported the notion that addition or promotion of individually or communally protective bacteria (such as probiotics) could minimize microbiota dysbiosis or directly prevent infection to restore host health (10–12).

Although the efficacy of probiotics has been shown in animal and humans, their mechanisms of action are poorly understood and low throughput experimental models often offer limited information on the individual contribution of probiotic species to community functions (1, 6, 7, 13, 14). Moreover, characterization of bacterial strains improving colonization resistance is still hindered by the complexity of host-commensal ecosystems. Zebrafish have recently emerged as a powerful tool to study microbe-microbe and host-microbe interactions (15–19). Zebrafish can be easily reared germ-free or gnotobiotically in association with specific bacterial species (15, 20). Moreover, zebrafish bacterial communities are increasingly well characterized and a number of phylogenetically distinct zebrafish gut bacteria can be cultured, making this model system directly amenable to microbiota manipulation and assessment of probiotic effect on host infection resistance (21–24). Several studies have used zebrafish to evaluate the effect of exogenous addition of potential probiotics on host resistance to infection by various pathogens (22–29). However, the role of the endogenous microbial community in protecting against invasive pathogen was rarely assessed and the reported protections were often partial, illustrating the difficulty in identifying fully protective exogenous probiotics.

Here we used germ-free and conventional zebrafish larvae to mine the indigenous commensal microbiota for bacterial species protecting against *Flavobacterium columnare*, a bacterial pathogen affecting wild and cultured fish species. We identified two distinct infection resistance strategies preventing mortality caused by *F. columnare*, mediated either by an individual member of the microbiota, the *Bacteroidetes Chryseobacterium massiliae* or by an assembly of 9 individually non-protecting bacterial species that formed a protective community. Our results demonstrated that mining host microbiota constitutes a powerful approach to identify key mediators of intrinsic colonization resistance, providing insight into how to engineer ecologically resilient and protective microbial communities.

## MATERIALS AND METHODS

### Bacterial strains and growth conditions

Bacterial strains isolated from zebrafish microbiota are listed in Table 1. *F. columnare* strains (Suppl. Table S3) were grown at 28°C in tryptone yeast extract salts (TYES) broth [0.4 % (w/v) tryptone, 0.04 % yeast extract, 0.05 % (w/v) MgSO_4_ 7H_2_O, 0.02 % (w/v) CaCl_2_ 2H_2_O, 0.05 % (w/v) D-glucose, pH 7.2]. *F. columnare* strains were assigned into four genomovar groups using 16S rRNA restriction fragment length polymorphism analysis, including genomovar I, I/II, II, and III (30). All 10 strains of the core zebrafish microbiota species were grown in TYES or LB at 28°C.

**Table 1.** The 10 strains composing the core of zebrafish larvae microbiota. Bacterial strains consistently detected at all time points (6 and 11 dpf) in all experiment runs and constituting the core of conventional zebrafish larval microbiota and their taxonomic affiliation.

| Bacterial species of the core zebrafish microbiota | ANI %<sup>a</sup> | 16S rRNA (%)<sup>b</sup> | *recA* (%)<sup>c</sup> | *rplC* (%)<sup>d</sup> |
| --- | --- | --- | --- | --- |
| *Aeromonas veronii* 1 | 96.52 | 98.27 | 97.00 | 99.84 |
| *Aeromonas veronii* 2 | 96.58 | 99.53 | 98.31 | 99.68 |
| *Aeromonas caviae* | 97.97 | 99.94 | 98.78 | 99.84 |
| *Chryseobacterium massiliae* | 95.85 | 99.86 | 96.61 | 99.84 |
| *Phyllobacterium myrsinacearum* | 98.58 | 99.86 | 99.72 | 100 |
| *Pseudomonas sediminis* | 96.12 | 99.73 | 97.70 | 99.84 |
| *Pseudomonas mossellii* | 99.39 | 98.27 | 100 | 99.84 |
| *Pseudomonas nitroreducens* * | 92.14 | 99.80 | 94.95 | 99.06 |
| *Pseudomonas peli* * | 88.84 | 99.20 | 91.51 | 95.44 |
| *Stenotrophomas maltophilia* * | 90.94 | 97.85 | 95.38 | 99.08 |

<sup>a</sup> Average Nucleotide Identity value
<sup>b</sup> 16S rRNA gene sequence similarity
<sup>c</sup> *recA* gene sequence similarity
<sup>d</sup> *rplC* gene sequence similarity
\*Species ambiguously identified

### Ethics statement

All animal experiments described in the present study were conducted at the Institut Pasteur (larvae) or at INRA Jouy-en-Josas (adults) according to European Union guidelines for handling of laboratory animals (http://ec.europa.eu/environment/chemicals/lab_animals/home_en.htm) and were approved by the relevant institutional Animal Health and Care Committees.

### General handling of zebrafish

Wild-type AB fish, originally purchased from the Zebrafish International Resource Center (Eugene, OR, USA), or *myd88*-null mutants (*myd88*^*hu3568/hu3568*^) (31), kindly provided by A.H Meijer, (Leiden University, the Netherlands), were raised in our facility. A few hours after spawning, eggs were collected, rinsed, and sorted under a dissecting scope to remove faeces and unfertilized eggs. All following procedures were performed in a laminar microbiological cabinet with single-use disposable plasticware. Fish were kept in sterile 25 cm^3^ vented cap culture flasks containing 20 mL of water (0-6 dpf-15 fish per flasks) or 24-well microtiter plates (6-15 dpf-1 fish per 2 mL well) in autoclaved mineral water (Volvic) at 28°C. Fish were fed 3 times a week from 4 dpf with germ-free *Tetrahymena thermophila* protozoans (22). Germ-free zebrafish were produced after sterilizing the egg chorion protecting the otherwise sterile egg, with antibiotic and chemical treatments (see below), whereas conventional larvae (with facility-innate microbiota) were directly reared from non-sterilized eggs and then handled exactly as the germ-free larvae.

### Sterilization of zebrafish eggs

Egg sterilization was performed as previously described with some modifications (22). Freshly fertilized zebrafish eggs were first bleached (0.003%) for 5 min, washed 3 times in sterile water under gentle agitation and maintained overnight in groups of 100 eggs per 75 cm^3^ culture flasks with vented caps containing 100 mL of autoclaved Volvic mineral water supplemented with methylene blue solution (0.3 μg/mL). Afterwards, eggs were transferred into 50 mL Falcon tubes (100 eggs per tube) and treated with a mixture of antibiotics (500 μL of penicillin G: streptomycin, 10,000 U/ml: 10 mg/mL GIBCO #P4333), 200 μL of filtered kanamycin sulfate (100 mg/mL, SERVA Electrophoresis #26899) and antifungal drug (50 μL of amphotericin B solution Sigma-Aldrich (250 μg/mL) #A2942) for 2 h under agitation at 28°C. Eggs were then washed 3 times in sterile water under gentle agitation and bleached (0.003%) for 15 min, resuspending the eggs every 3 min by inversion. Eggs were washed again 3 times in water and incubated 10 min with 0.01% Romeiod (COFA, Coopérative Française de l’Aquaculture). Finally, eggs were washed 3 times in water and transferred into 25 cm^3^ culture flasks with vented caps containing 20 mL of water. After sterilization, eggs were transferred with approximately 30 to 35 eggs / flasks and were transferred into new flasks at 4 dpf before re-conventionalization with 10 to 15 fish / flask. We monitored sterility at several points during the experiment by spotting 50 μL of water from each flask on LB, TYES and on YPD agar plates, all incubated at 28°C under aerobic conditions. Plates were left for at least 3 days to allow slow-growing organisms to multiply. Spot checks for bacterial contamination were also carried out by PCR amplification of water samples with the 16S rRNA gene primers and procedure detailed further below. If a particular flask was contaminated, those fish were removed from the experiment.

### Procedure for raising germ-free zebrafish

After hatching, fish were fed with germ-free *T. thermophila* 3 times per week from 4 dpf onwards. *(i) T. thermophila stock.* A germ-free line of *T. thermophila* was maintained at 28°C in 20 mL of PPYE (0.25% proteose peptone BD Bacto #211684, 0.25% yeast extract BD Bacto #212750) supplemented with penicillin G (10 unit/mL) and streptomycin (10μg/mL). Medium was inoculated with 100 μL of the preceding *T. thermophila* stock. After one week of growth, samples were taken, tested for sterility on LB, TYES and YPD plates and restocked again. *(ii) Growth. T. thermophila* were incubated at 28°C in MYE broth (1% milk powder, 1% yeast extract) inoculated from stock suspension at a 1:50 ratio. After 24 h of growth, *T. thermophila* were transferred to Falcon tubes and washed (4400 rpm, 3 min at 25°C) 3 times in 50 mL of autoclaved Volvic water. Finally, *T. thermophila* were resuspended in sterile water and added to culture flasks (500 μL in 20 mL) or 24-well plates (50 μL / well). Sterility of *T. thermophila* was tested by plating and 16S rRNA PCR as described in the section above. (iii) *Fine-powder feeding*. When indicated, fish were fed with previously γ-ray-sterilized fine-powdered food suitable for an early first feeding gape size (ZM-000 fish feed, ZM Ltd) every 48 hours (32).

### >Re-conventionalization of germ-free zebrafish

At 4 dpf, just after hatching, zebrafish larvae were re-conventionalized with a single bacterial population or a mix of several. The 10 bacterial strains constituting the core protective microbiota were grown for 24h in suitable media (TYES or LB) at 28°C. Bacteria were then pelleted and washed twice in sterile water, and all adjusted to the same cell density (OD_600_ = 1 or 5.10^7^ cfu/mL) (i) *Re-conventionalization with individual species.* Bacteria were resuspended and transferred to culture flasks containing germ-free fish at a final concentration of 5.10^5^ cfu/mL. (ii) *Re-conventionalization with bacterial mixtures*. For the preparation of Mix10, Mix9, Mix8 and all other mixes used, equimolar mixtures were prepared by adding each bacterial species at initial concentration to 5.10^7^ cfu/mL. Each bacterial mixture suspension was added to culture flasks containing germ-free fish at a final concentration of 5.10^5^ cfu/mL.

### Infection challenges

*F. columnare* strains (Suppl. Table S3) were grown overnight in TYES broth at 28°C. Then, 2 mL of culture were pelleted (10,000 rpm for 5 min) and washed once in sterile water. GF zebrafish were brought in contact with the tested pathogens at 6 dpf for 3h by immersion in culture flasks with bacterial doses ranging from 5.10^2^ to 5.10^7^ cfu/mL. Fish were then transferred to individual wells of 24-well plates, containing 2 mL of water and 50 μL of freshly prepared GF *T. thermophila* per well. Mortality was monitored daily as described in (22) and as few as 54±9 cfu/larva of *F. columnare* were recovered from infected larvae. All zebrafish experiments were stopped at day 9 post-infection and zebrafish were euthanized with tricaine (MS-222) (Sigma-Aldrich #E10521). Each experiment was repeated at least 3 times and between 10 and 15 larvae were used per condition and per experiment.

### Collection of eggs from other zebrafish facilities

Conventional zebrafish eggs were collected in 50 mL Falcon tubes from the following facilities: Facility 1 - zebrafish facility in Hospital Robert Debré, Paris; Facility 2 - Jussieu zebrafish facility A2, University Paris 6; Facility 3 - Jussieu – zebrafish facility C8 (UMR7622), University Paris 6; Facility 4: AMAGEN commercial facility, Gif sur Yvette; Larvae were treated with the same rearing conditions, sterilization and infection procedures used in the Institut Pasteur facility.

### Determination of fish bacterial load using cfu count

Zebrafish were euthanized with tricaine (MS-222) (Sigma-Aldrich #E10521) at 0.3 mg/mL for 10 minutes. Then they were washed in 3 different baths of sterile PBS-0.1% Tween to remove bacteria loosely attached to the skin. Finally, they were transferred to tubes containing calibrated glass beads (acid-washed, 425 μm to 600 μm, SIGMA-ALDRICH #G8772) and 500 μL of autoclaved PBS. They were homogenized using FastPrep Cell Disrupter (BIO101/FP120 QBioGene) for 45 s at maximum speed (6.5 m/s). Finally, serial dilutions of recovered suspension were spotted on TYES agar and cfu were counted after 48h of incubation at 28°C.

### Characterization of zebrafish microbial content

Over 3 months, the experiment was run independently 3 times and 3 different batches of eggs were collected from different fish couples in different tanks. Larvae were reared as described above. GF and Conv larvae were collected at 6 dpf and 11 dpf for each batch. Infected Conv larvae were exposed to *F. columnare*^ALG^ for 3h by immersion as described above. For each experimental group, triplicate pools of 10 larvae (one in each experimental batch) were euthanized, washed and lysed as above. Lysates were split into 3 aliquots, one for culture followed by 16S rRNA gene sequencing (A), one for 16S rRNA gene clone library generation and Sanger sequencing (B), and one for Illumina metabarcoding-based sequencing (C).

#### A) Bacterial culture followed by 16S rRNA gene-based identification

Lysates were serially diluted and immediately plated on R2A, TYES, LB, MacConkey, BHI, BCYE, TCBS and TSB agars and incubated at 28°C for 24-72h. For each agar, colony morphotypes were documented, and colonies were picked and re-streaked on the same agar in duplicate. In order to identify the individual morphotypes, individual colonies were picked for each identified morphotype from each agar, vortexed in 200 μL DNA-free water and boiled for 20 min at 90°C. Five μL of this bacterial suspension were used as template for colony PCR to amplify the 16S rRNA gene with the universal primer pair for the Domain bacteria 8f (5’-AGA GTT TGA TCC TGG CTC AG-3’) and 1492r (5’-GGT TAC CTT GTT ACG ACT T-3’). Each primer was used at a final concentration of 0.2 μM in 50 μL reactions. PCR cycling conditions were - initial denaturation at 94°C for 2 min, followed by 32 cycles of denaturation at 94 °C for 1 min, annealing at 56°C for 1 min, and extension at 72°C for 2 min, with a final extension step at 72°C for 10 min. 16S rRNA gene PCR products were verified on 1% agarose gels, purified with the QIAquick® PCR purification kit and two PCR products for each morphotype were sent for sequencing (Eurofins, Ebersberg, Germany). 16S rRNA sequences were manually proofread, and sequences of low quality were removed from the analysis. Primer sequences were trimmed, and sequences were compared to GenBank (NCBI) with BLAST, and to the Ribosomal Database Project with SeqMatch. For genus determination a 95% similarity cut-off was used, for Operational Taxonomic Unit determination, a 98% cut-off was used.

#### B) 16S rRNA gene clone library generation

Total DNA was extracted from the lysates with the Mobio PowerLyzer® Ultraclean® kit according to manufacturer’s instructions. Germ-free larvae and DNA-free water were also extracted as control samples. Extracted genomic DNA was verified by Tris-acetate-EDTA-agarose gel electrophoresis (1%) stained with GelRed and quantified by applying 2.5 μL directly to a NanoDrop^®^ ND-1000 Spectrophotometer. The 16S rRNA gene was amplified by PCR with the primers 8f and 1492r, and products checked and purified as described in section A. Here, we added 25-50 ng of DNA as template to 50 μL reactions. Clone libraries were generated with the pGEM®-T Easy Vector system (Promega) according to manufacturer’s instructions. Presence of the cloned insert was confirmed by colony PCR with vector primers gemsp6 (5’-GCT GCG ACT TCA CTA GTG AT-3’) and gemt7 (5’-GTG GCA GCG GGA ATT CGA T-3’). Clones with an insert of the correct size were purified as above and sent for sequencing (Eurofins, Ebersberg, Germany). Blanks using DNA-free water as template were run for all procedures as controls. For the three independent runs of the experiment, 10 Conv fish per condition (6 and 11 dpf, exposed or not to *F. columnare*) and per repeat were pooled. Each pool of 10 fish was sequenced separately, generating 3 replicates for each condition (n=12), resulting in a total of 857 clones. Clone library coverage was calculated with the following formula [1-(n1/N2)] × 100, where n1 is the number of singletons detected in the clone library, and N_2_ is the total number of clones generated for this sample. Clone libraries were generated to a minimum coverage of 95%. Sequence analysis and identification was carried out as in section A.

#### C) by 16S rRNA V3V4 Amplicon Illumina sequencing

To identify the 16S rRNA gene diversity in our facility and fish collected from 4 other zebrafish facilities, fish were reared as described above. GF fish were sterilised as above, and uninfected germ-free and conventional fish were collected at 6 dpf and 11 dpf. Infection was carried out as above with *F. columnare*^ALG^ for 3h by bath immersion, followed by transfer to clean water. Infected conventional fish were collected at 6 dpf 6h after infection with *F. columnare* and at 11 dpf, the same as uninfected fish. GF infected larvae were only collected at 6 dpf 6h post infection, as at 11 dpf all larvae had succumbed to infection. Triplicate pools of 10 larvae were euthanized, washed and lysed as above. Total DNA was extracted with the Mobio PowerLyzer® Ultraclean® kit as described above and quantified with a NanoDrop^®^ ND-1000 Spectrophotometer and sent to IMGM Laboratories GmbH (Germany) for Illumina sequencing. Primers Bakt_341F (5’-CCTACGGGNGGCWGCAG-3’) and Bakt_805R (5’-GACTACHVGGGTATCTAATCC-3’), amplifying variable regions 3 and 4 of the 16S gene were used for amplification (33). Each amplicon was purified with solid phase reversible immobilization (SPRI) paramagnetic bead-based technology (AMPure XP beads, Beckman Coulter) with a Bead:DNA ratio of 0.7:1 (v/v) following manufacturer’s instructions. Amplicons were normalized with the Sequal-Prep Kit (Life Technologies), so each sample contained approximately 1 ng/μl DNA. Samples, positive and negative controls were generated in one library. The High Sensitivity DNA LabChip Kit (was used on the 2100 Bioanalyzer system (both Agilent Technologies) to check the quality of the purified amplicon library. For cluster generation and sequencing, MiSeq® reagents kit 500 cycles Nano v2 (Illumina Inc.) was used. Before sequencing, cluster generation by two-dimensional bridge amplification was performed, followed by bidirectional sequencing, producing 2 × 250 bp paired-end (PE) reads.

MiSeq® Reporter 2.5.1.3 software was used for primary data analysis (signal processing, de-multiplexing, trimming of adapter sequences). CLC Genomics Workbench 8.5.1 (Qiagen) was used for read-merging, quality trimming and QC reports and OTU definition were carried out in the CLC plugin Microbial Genomics module.

### Comparison of whole larvae vs intestinal bacterial content

Larvae re-conventionalized with Mix10 and infected with *F. columnare*^ALG^ at 6 dpf for 3h were euthanized and washed. DNA was extracted from pools of 10 whole larvae or of pools of 10 intestinal tubes dissected with sterile surgical tweezer and subjected to Illumina 16S rRNA gene sequencing. GF larvae and dissected GF intestines were sampled as controls. As dissection of the larval guts involved high animal loss and was a potential important contamination source, we proceeded with using entire larvae for the rest of the study.

### Whole genome sequencing

Chromosomal DNA of the ten species composing the core of zebrafish larvae microbiota was extracted using the DNeasy Blood & Tissue kit (QIAGEN) including RNase treatment. DNA quality and quantity were assessed on a NanoDrop ND-1000 spectrophotometer (Thermo Scientific).

DNA sequencing libraries were made using the Nextera DNA Library Preparation Kit (Illumina Inc.) and library quality was checked using the High Sensitivity DNA LabChip Kit on the Bioanalyzer 2100 (Agilent Technologies). Sequencing clusters were generated using the MiSeq reagents kit v2 500 cycles (Illumina Inc.) according to manufacturer’s instructions. DNA was sequenced at the Helmholtz Centre for Infection Research by bidirectional sequencing, producing 2 × 250 bp paired-end (PE) reads. Between 1,108,578 and 2,914,480 reads per sample were retrieved with a median of 1,528,402. Reads were quality filtered, trimmed and adapters removed with trimmomatic 0.39 (34) and genomes assembled using SPAdes 3.14 (35).

### Bacterial species identification

Whole genome-based bacterial species identification was performed by the TrueBac ID system (v1.92, DB:20190603) [https://www.truebacid.com/; (36). Species-level identification was performed based on the algorithmic cut-off set at 95% ANI when possible or when the 16S rRNA gene sequence similarity was >99 %.

### Monitoring of bacterial dynamics

Three independent experiments were run over 6 weeks with eggs collected from different fish couples from different tanks to monitor establishment and recovery. Larvae were reared, sterilized and infected as above with the only difference that 75 cm^3^ culture flasks with vented caps (filled with 50 mL of sterile Volvic) were used to accommodate the larger number of larvae required, as in each experiment. Larvae for time course Illumina sequencing were removed sequentially from the experiment that monitored the survival of the larvae. Animals were pooled (10 larvae for each time point/condition), euthanized, washed and lysed as described above and stored at −20°C until the end of the survival monitoring, and until all triplicates had been collected.

#### A) Community establishment

In order to follow the establishment of the 10 core strains in the larvae, GF larvae were re-conventionalized with an equiratio Mix10 as above. Re-con^Mix10^ larvae were sampled at 4 dpf immediately after addition of the 10 core species and then 20 min, 2h, 4h and 8h after. Germ-free, conventional larvae and the inoculum were also sampled as controls.

#### B) Induction of dysbiosis

Different doses of kanamycin (dose 1= 200 μg/mL; dose 2= 50 μg/mL; dose 3= 25 μg/mL) and a penicillin/streptomycin antibiotic mix (dose 1= 250 μg/mL; dose 2= 15.6 μg/mL were tested on re-con^Mix10^ 4 dpf zebrafish larvae by adding them to the flask water to identify antibiotic treatments that were non-toxic to larvae but that caused dysbiosis.

After 16 hours of treatment, antibiotics were extensively washed off with sterile water and larvae were challenged with *F. columnare*^ALG^, leading to the death of all larvae – e.g. successful abolition of colonization resistance with best results in all repeats with 250 μg/mL penicillin/streptomycin and 50 μg/mL kanamycin as antibiotic treatment.

#### C) Community recovery

As in B) after 8h of incubation, 4 dpf recon^Mix10^ larvae were treated with 250 μg/mL penicillin/streptomycin and 50 μg/mL kanamycin for 16h. Antibiotics were extensively washed off and larvae were now left to recover in sterile water for 24h to assess resilience of the bacterial community. Samples (pools of 10 larvae) were taken at 3h, 6h, 12h, 18h and 24h during recovery and sent for 16S rRNA Illumina sequencing. Larvae were then challenged at 6 dpf with *F. columnare*^ALG^ for 3h and survival was monitored daily for 9 days post-infection. All time course samples were sequenced by IMGM Laboratories GmbH, as described above.

### Statistical analysis of metataxonomic data

16S RNA analysis was performed with SHAMAN (37)]. Library adapters, primer sequences, and base pairs occurring at 5’ and 3’ends with a Phred quality score <20 were trimmed off by using Alientrimmer (v0.4.0). Reads with a positive match against zebrafish genome (mm10) were removed. Filtered high-quality reads were merged into amplicons with Flash (v1.2.11). Resulting amplicons were clustered into operational taxonomic units (OTU) with VSEARCH (v2.3.4) (38). The process includes several steps for de-replication, singletons removal, and chimera detection. The clustering was performed at 97% sequence identity threshold, producing 459 OTUs. The OTU taxonomic annotation was performed against the SILVA SSU (v132) database (39) completed with 16S sequence of 10 bacterial communities using VSEARCH and filtered according to their identity with the reference (40). Annotations were kept when the identity between the OTU sequence and reference sequence is ≥ 78.5% for taxonomic Classes, ≥ 82% for Orders, ≥ 86.5% for Families, ≥ 94.5% for Genera and ≥ 98% for species. Here, 73.2% of the OTUs set was annotated and 91.69 % of them were annotated at genus level.

The input amplicons were then aligned against the OTU set to get an OTU contingency table containing the number of amplicon associated with each OTU using VSEARCH global alignment. The matrix of OTU count data was normalized for library size at the OTU level using a weighted non-null count normalization. Normalized counts were then summed within genera. The generalized linear model (GLM) implemented in the DESeq2 R package (41) was then applied to detect differences in abundance of genera between each group. We defined a GLM that included the treatment (condition) and the time (variable) as main effects and an interaction between the treatment and the time. Resulting P values were adjusted according to the Benjamini and Hochberg procedure.

The statistical analysis can be reproduced on shaman by loading the count table, the taxonomic results with the target and contrast files that are available on figshare https://doi.org/10.6084/m9.figshare.11417082.v2.

### Determination of cytokine levels

Total RNAs from individual zebrafish larvae were extracted using RNeasy kit (Qiagen), 18h post pathogen exposure (12h post-wash). Oligo(dT17)-primed reverse transcriptions were carried out using M-MLV H-reverse-transcriptase (Promega). Quantitative PCRs were performed using Takyon SYBR Green PCR Mastermix (Eurogentec) on a StepOne thermocycler (Applied Biosystems). Primers for *ef1a* (housekeeping gene, used for cDNA amount normalization), *il1b*, *il10* and *il22* are described in (22). Data were analysed using the ΔΔCt method. Four larvae were analysed per condition. Zebrafish genes and proteins mentioned in the text: *ef1a* NM_131263*; il1b* BC098597*; il22* NM_001020792*; il10* NM_001020785*; myd88* NM_212814.

### Histological comparisons of GF, Conv and Re-Conv fish GF infected or not with *F. columnare*

Fish were collected 24h after infection (7 dpf) and were fixed for 24h at 4°C in Trump fixative (4% methanol-free formaldehyde, 1% glutaraldehyde in 0.1 M PBS, pH 7.2) and sent to the PIBiSA Microscopy facility services (https://microscopies.med.univ-tours.fr/) in the Faculté de Médecine de Tours (France), where whole fixed animals were processed, embedded in Epon. Semi-thin sections (1μm) and cut using a X ultra-microtome and then either dyed with toluidine blue for observation by light microscopy and imaging or processed for Transmission electron microscopy.

### Adult zebrafish pre-treatment with *C. massiliae*

The zebrafish line AB was used. Fish were reared at 28°C in dechlorinated recirculated water, then transferred into continuous flow aquaria when aging 3-4 months for infection experiments. *C. massiliae* was grown in TYES broth at 150 rpm and 28°C until stationary phase. This bacterial culture was washed twice in sterile water and adjusted to OD600nm = 1. Adult fish re-conventionalization was performed by adding *C. massiliae* bacterial suspension directly into the fish water (1L) at a final concentration of 2.10^5^ cfu/mL. Bacteria were maintained in contact with fish for 24 h by stopping the water flow then subsequently removed by restoring the water flow. *C. massiliae* administration was performed twice after water renewal. In the control group, the same volume of sterile water was added.

### Adult zebrafish infection challenge

*F. columnare* infection was performed just after fish re-conventionalization with *C. massiliae*. The infection was performed as previously described by Li and co-workers with few modifications [Li et al., 2017]. Briefly, *F. columnare* strain ALG-0530 was grown in TYES broth at 150 rpm and 28 °C until late-exponential phase. Then, bacterial cultures were diluted directly into the water of aquaria (200 mL) at a final concentration of 5.106 cfu/mL. Bacteria were maintained in contact with fish for 1 h by stopping the water flow then subsequently removed by restoring the water flow. Sterile TYES broth was used for the control group. Bacterial counts were determined at the beginning of the immersion challenge by plating serial dilutions of water samples on TYES agar. Water was maintained at 28°C and under continuous oxygenation for the duration of the immersion. Groups were composed of 10 fish. Virulence was evaluated according to fish mortality 10 days post-infection.

### Statistical methods

Statistical analyses were performed using unpaired, non-parametric Mann-Whitney test or unpaired t-tests. Analyses were performed using Prism v8.2 (GraphPad Software).

Evenness: The Shannon diversity index was calculated with the formula (HS = −Σ[P(ln(P)]) where P is the relative species abundance. Total evenness was calculated for the Shannon index as E = H_S_/H_max_. The less evenness in communities between the species (and the presence of a dominant species), the lower this index is.

### Data availability

Bacterial genome sequences obtained in the present study are available at the European Nucleotide Archive with the project number PRJEB36872, under accession numbers numbers ERS4385993 (*Aeromonas veronii 1*); ERS4386000 (*Aeromonas veronii 2*); ERS4385996 (*Aeromonas caviae*); ERS4385998 (*Chryseobacterium massiliae*); ERS4385999 (*Phyllobacterium myrsinacearum*); ERS4406247 (*Pseudomonas sediminis*); ERS4385994 (*Pseudomonas mossellii*) ERS4386001 (*Pseudomonas nitroreducens*); ERS4385997 (*Pseudomonas peli*); ERS4385995 (*Stenotrophomas maltophilia*);

## RESULTS

### *Flavobacterium columnare* kills germ-free but not conventional zebrafish

To investigate microbiota-based resistance to infection in zebrafish, we compared the sensitivity of germ-free (GF) and conventional (Conv) zebrafish larvae to *F. columnare*, an important fish pathogen affecting carp, channel catfish, goldfish, eel, salmonids and tilapia and previously shown to infect and kill adult zebrafish (12, 42–45). We used bath immersion to expose GF and Conv zebrafish larvae at 6 days post fertilization (dpf), to a collection of 28 *F. columnare* strains, belonging to four different genomovars for 3h at 5.10^5^ colony forming units (cfu)/mL. Daily monitoring showed that 16 out of 28 *F. columnare* strains killed GF larvae in less than 48h (Supplementary Fig. S1A), whereas Conv larvae survived exposure to all tested virulent *F. columnare* strains (Supplementary Fig. S1B). Exposure to the highly virulent strain ALG-00-530 (hereafter *F. columnare*^ALG^) also showed that GF mortality was fast (1 day) and dose-dependent and that Conv zebrafish survived all but the highest dose (10^7^ cfu/mL) (Fig. 1). Similar survival of infected Conv larvae was obtained with zebrafish AB strain eggs obtained from 4 different zebrafish facilities (Supplementary Fig. S2), suggesting that conventional zebrafish microbiota could provide protection against *F. columnare* infection.

**Figure 1.**
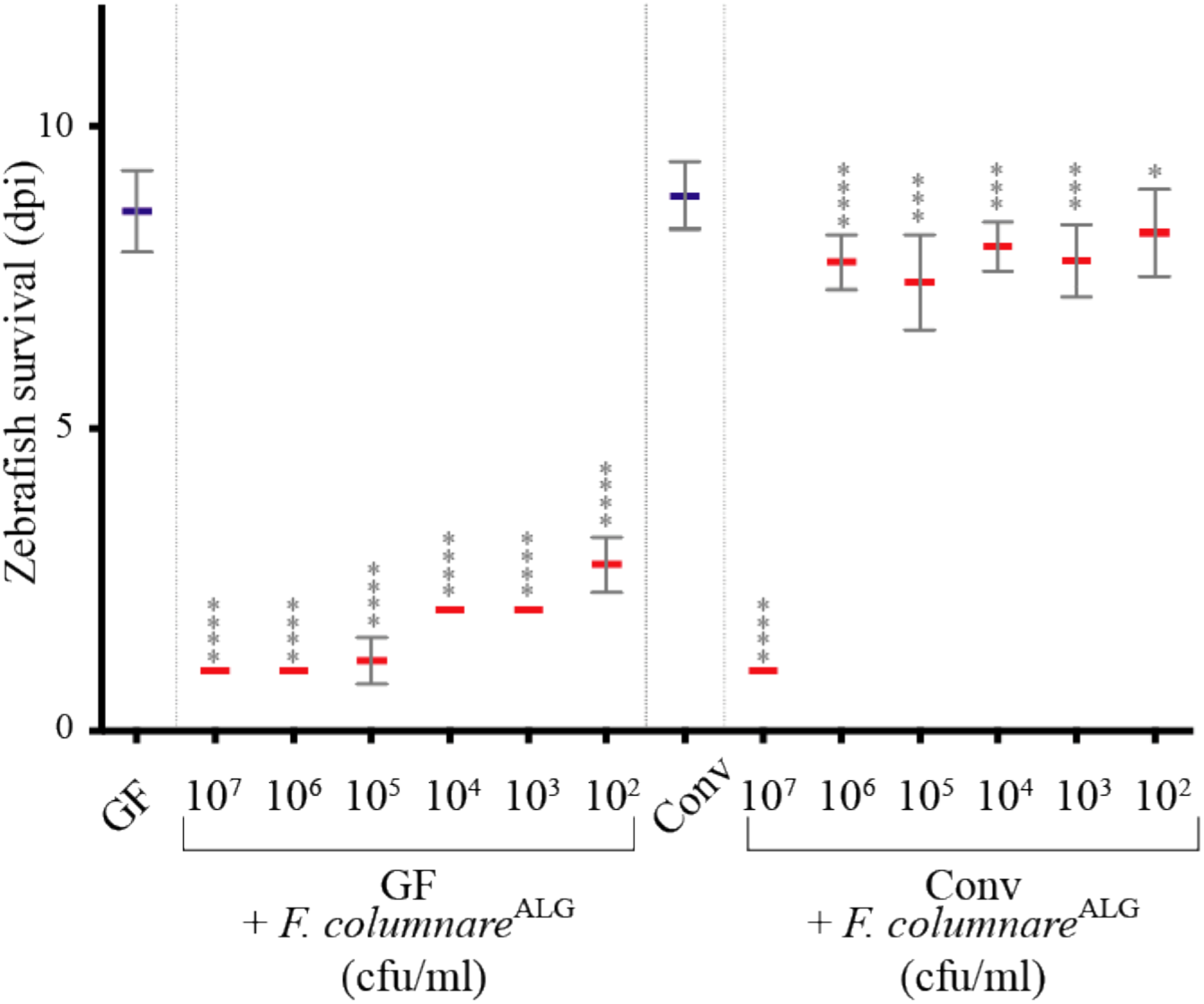
*Flavobacterium columnare* kills germ-free but not conventional zebrafish. 6 dpf (corresponding to 0 dpi) GF or Conv zebrafish larvae were exposed to different doses of *F. columnare*^ALG^ by bath immersion and transferred after 3h into sterile water. Mean survival is represented by a thick horizontal bar with standard deviation. For each condition, n = 12 zebrafish larvae. Larvae mortality rate was monitored daily and surviving fish were euthanized at day 9 post infection. Statistics correspond to unpaired, non-parametric Mann-Whitney test comparing all conditions to non-infected GF (left) or Conv (right). ****: p<0.0001; ***: p<0.001; *: p<0.05, absence of *: non-significant. Blue mean bars correspond to non-exposed larvae and red mean bars correspond to larvae exposed to *F. columnare*.

### A community of 10 culturable bacterial strains protect against *F. columnare* infection

In our rearing conditions, the conventional larval microbiota is acquired after hatching from microorganisms present on the egg chorion and in fish facility water. To test the hypothesis that microorganisms associated with conventional eggs provided protection against *F. columnare*^ALG^, we exposed sterilized eggs to either fish facility tank water or to non-sterilized conventional eggs at 0 or 4 dpf (before or after hatching, respectively). In both cases, these re-conventionalized (re-Conv) zebrafish survived *F. columnare*^ALG^ infection as well as Conv zebrafish (Supplementary Fig. S3). To determine the composition of conventional zebrafish microbiota, we generated 16S rRNA gene clone libraries from homogenate pools of Conv larvae aged 6 and 11 dpf exposed or not to *F. columnare*^ALG^, sampled over 3 months from 3 different batches of larvae (n=10). A total of 857 clones were generated for all samples. We identified 15 operational taxonomical units (OTUs), 10 of which were identified in all experiments (Table 1, Suppl. Table S1). Two OTUs (belonging to an *Ensifer* sp. and a *Hydrogenophaga* sp.) were only detected once, and a *Delftia* sp., a *Limnobacter* sp. and a *Novosphingobium* sp. were detected more than once (2, 3 and 2 times, respectively), but not consistently in all batches of fish (Table 1, Suppl. Table S1). Moreover, deep-sequencing of the 16S rRNA V3-V4 region of gDNA retrieved from larvae originating from the other four zebrafish facilities described above, revealed that OTUs for all of these 10 strains were also detected in Conv larvae, with the exception of *A. veronii* strain 2 that was not detected in all samples (Suppl. Table S2).

To isolate culturable zebrafish microbiota bacteria, we plated dilutions of homogenized 6 dpf and 11 dpf larvae pools on various growth media and we identified 10 different bacterial morphotypes. 16S-based analysis followed by full genome sequencing identified 10 bacteria corresponding to 10 strains of 9 different species that were also consistently detected by culture-free approaches (Table 1). To assess the potential protective role of these 10 strains, we re-conventionalized GF zebrafish at 4 dpf with a mix of all 10 identified culturable bacterial species (hereafter called Mix10), each at a concentration of 5.10^5^ cfu/mL and we monitored zebrafish survival after exposure to *F. columnare*^ALG^ at 6 dpf. We showed that zebrafish re-conventionalized with the Mix10 (Re-Conv^Mix10^) displayed a strong level of protection against all identified highly virulent *F. columnare* strains (Supplementary Fig. S4). These results demonstrated that the Mix10 constitutes a core protective bacterial community providing full protection of zebrafish larvae against *F. columnare* infection.

### Community dynamics under antibiotic-induced dysbiosis reveal a key contributor to resistance to *F. columnare* infection

To further analyze the determinants of Mix10 protection against *F. columnare*^ALG^ infection, we inoculated 4 dpf larvae with an equal-ratio mix of the 10 bacteria (at 5.10^5^ cfu/mL each) and monitored their establishment over 8 hours. We first verified that whole larvae bacterial content (OTU abundance) was not significantly different from content of dissected intestinal tubes (p=0.99, two-tailed t-test) (Supplementary Fig. S5) and proceeded to use entire larvae to monitor bacterial establishment and recovery in the rest of the study. We then collected pools of 10 larvae immediately after re-conventionalization (t0), and then at 20 min, 2 hours, 4 hours and 8 hours in three independent experiments. Illumina sequencing of the 16S rRNA gene was used to follow bacterial relative abundance. At t0, all species were present at > 4% in the zebrafish, apart from *A. veronii* strains 1 (0.2%) and 2 (not detected) (Supplementary Fig. S6). *Aeromonas caviae* was detected as the most abundant species (33%), followed by *Stenotrophomonas maltophilia* (23%) and *Chryseobacterium massiliae* (12%), altogether composing 68% of the community (Supplementary Fig. S6). The relative species abundance, possibly reflecting initial colonization ability, was relatively stable for most species during community establishment, with similar species evenness at t0 (E = 0.84) and t8h (E = 0.85). Whereas both Conv and Re-Conv^Mix10^ larvae were protected against *F. columnare*^ALG^ infection, the global structure of the reconstituted Mix10 population was different from the conventional one at 4 dpf (Supplementary Fig. S6).

To test the sensitivity to disturbance and the resilience of the protection provided by Mix10 bacterial community, we subjected Re-Conv^Mix10^ zebrafish to non-toxic antibiotic treatment at 4dpf using either 250 μg/mL penicillin/streptomycin combination (all members of the Mix10 bacteria are sensitive to penicillin/streptomycin) or 50 μg/mL kanamycin (affecting all members of the Mix10 bacteria except *C. massiliae*, *P. myrsinacearum* and *S. maltophilia*) (Suppl. Fig. S7). At 5 dpf, after 16 hours of exposure, antibiotics were washed off and zebrafish were immediately exposed to *F. columnare*^ALG^. Both antibiotic treatments resulted in complete loss of the protection against *F. columnare*^ALG^ infection observed in Re-Conv^Mix10^ (Fig. 2A). We then used the same antibiotic treatments but followed by a 24h recovery period after washing off the antibiotics at 5 dpf, therefore only performing the infection at 6 dpf (Fig. 2B). Whilst Re-Conv^Mix10^ larvae treated with penicillin/streptomycin showed similar survival to infected GF larvae, kanamycin-treated Re-Conv^Mix10^ zebrafish displayed restored protection after 24h recovery and survived similarly to untreated conventionalized fish (Fig. 2B). Sampling and 16S gene analysis during recovery experiments at different time points showed that bacterial community evenness decreased after antibiotic administration for both treatments (E = 0.85 for 4 dpf control, E = 0.72 for t_0_ kanamycin and E = 0.7 for t_0_ penicillin/streptomycin), and continued to decrease during recovery (E = 0.6 and 0.64 for kanamycin and penicillin/streptomycin treatment after 24h recovery, respectively). Although *C. massiliae* remained detectable immediately after both antibiotic treatments, penicillin/streptomycin treatment led a significant reduction in its relative abundance (0.21%) (Fig. 2C). By contrast, *C. massiliae* relative abundance rebounded 6h after cessation of kanamycin treatment and was the dominant member (52%) of the reconstituted microbiota after 24h recovery period (Fig. 2D), suggesting that the protective effect observed in the kanamycin-treated larvae might be due to the recovery of *C. massiliae*.

**Figure 2.**
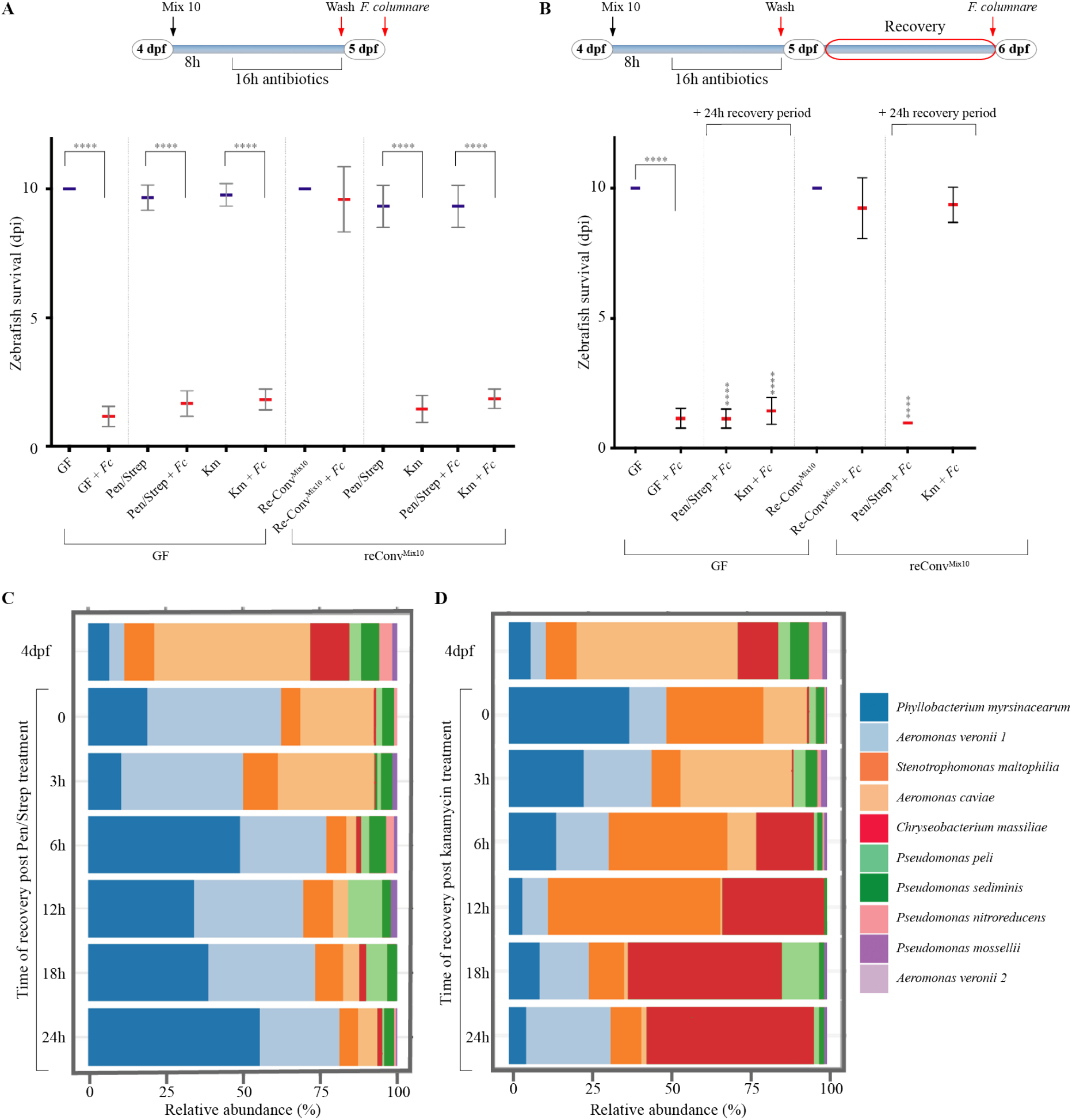
Analysis of protection against *F. columnare* infection after antibiotic dysbiosis. **A:** Response of zebrafish larvae to exposure to *F. columnare*^ALG^ after antibiotic-induced dysbiosis with a diagram showing timing and treatments of the experiment. **B:** A 24h period after antibiotic treatment allows the recovery of protection in kanamycin-treated zebrafish larvae with a diagram showing timing and treatments. Mean survival is represented by a thick horizontal bar with standard deviation. For each condition, n = 12 zebrafish larvae. Blue mean bars correspond to larvae not exposed to the pathogen and red mean bars correspond to exposed larvae. Larvae mortality rate was monitored daily and surviving fish were euthanized at day 9 post exposition to the pathogen. Indicated statistics correspond to unpaired, non-parametric Mann-Whitney test. ****: p<0.0001; absence of *: non-significant. **C:** Community recovery profile with streptomycin/penicillin treatment. **D:** Community recovery profile with kanamycin treatment. Pools of 10 larvae were collected for 16S rRNA sequencing.

### Resistance to *F. columnare* infection is provided by both individual and community-level protection

To test the potential key role played by *C. massiliae* in protection against *F. columnare*^ALG^ infection, we exposed 4 dpf GF zebrafish to *C. massiliae* only and showed that it conferred individual protection at doses as low as 5.10^2^ cfu/mL (Fig. 3). Whereas none of the 9 other species composing the Mix10 were protective individually (Fig. 3A), their equiratio combination (designated as Mix9) conferred protection to zebrafish, although not at doses lower than 5.10^4^ cfu/mL and not as reproducibly as with *C. massiliae* (Fig. 3B). To identify which association of species protected Re-Conv^Mix9^ zebrafish against *F. columnare*^ALG^ infection, we tested all 9 combinations of 8 species (Mix8), as well as several combinations of 7, 6, 4 or 3 species and showed no protection (Suppl. Fig. S8A and Suppl. Table S4). We then tested whether lack of protection of Mix8 compared to Mix9 could rely on a density effect by doubling the concentration of any of the species within the non-protective Mix8a (Suppl. Fig. S8B) and showed no protection. These results therefore indicated that microbiota-based protection against *F. columnare*^ALG^ infection relied on either *C. massiliae-*dependent membership effect or on a community-dependent effect mediated by the Mix9 consortium.

**Figure 3.**
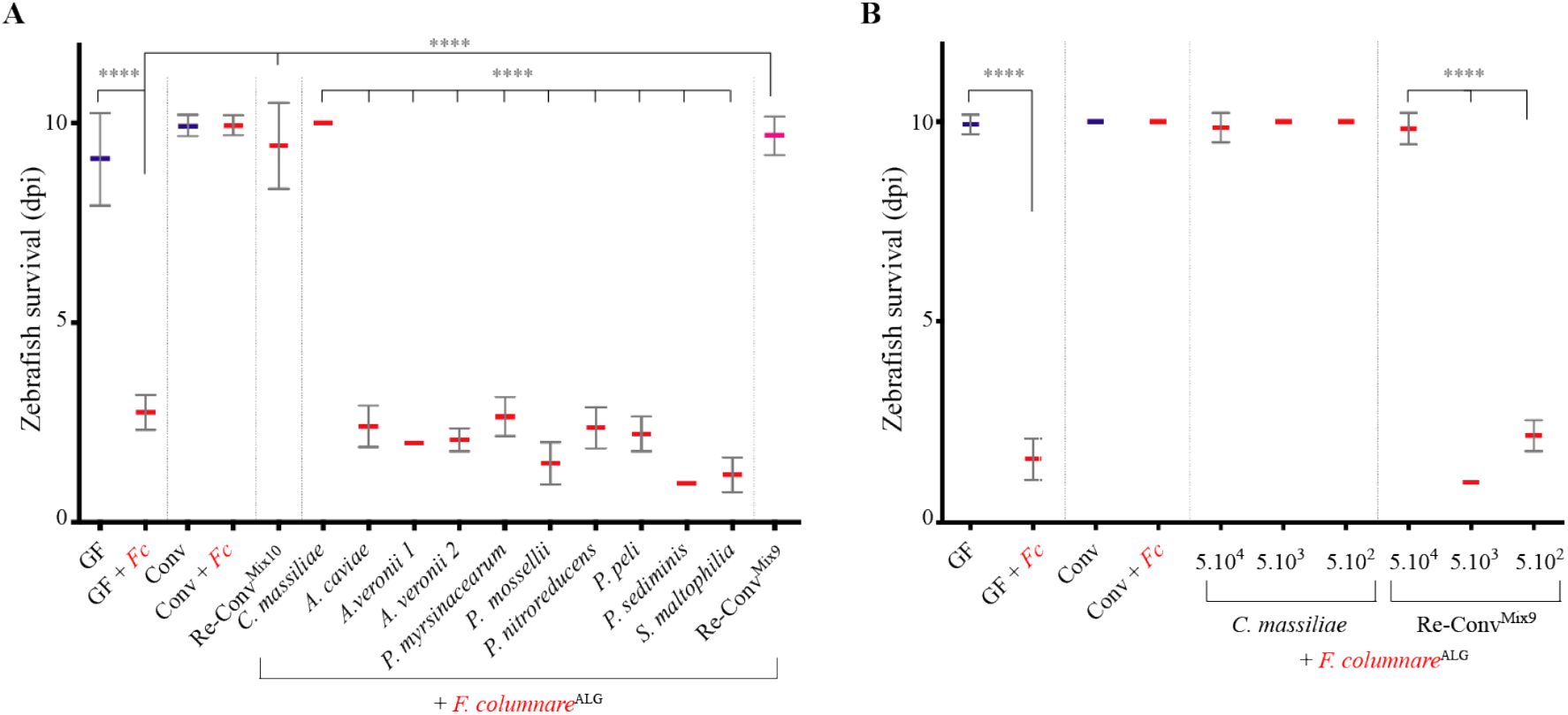
Protection against *F. columnare* in zebrafish re-conventionalized with individual or mixed bacterial strains isolated from zebrafish. **A**: Determination of the level of protection provided by each of the 10 bacterial strains composing the core protective zebrafish microbiota. Bacteria were added individually to the water on hatching day (dose 5.10^5^ cfu/mL). **B**: Level of protection provided by different amount of *C. massiliae* and Mix9. Mix9 only protected at the highest inoculum doses. Mean survival is represented by a thick horizontal bar with standard deviation. For each condition, n = 12 zebrafish larvae. Blue mean bars correspond to larvae not exposed to the pathogen and red mean bars correspond to exposed larvae. Larvae mortality rate was monitored daily and surviving fish were euthanized at day 9 post exposition to the pathogen. Indicated statistics correspond to unpaired, non-parametric Mann-Whitney test. ****: p<0.0001; absence of *: non-significant.

### Pro- and anti-inflammatory cytokine production does not contribute to microbiota-mediated protection against *F. columnare*^ALG^ infection

To test the contribution of the immune response of zebrafish larvae to resistance to *F. columnare* infection, we used qRT-PCR to measure cytokine mRNA expression in GF and Conv zebrafish exposed or not to *F. columnare*^ALG^. We also tested the impact of re-conventionalization with *C. massiliae* (re-Conv^*Cm*^), Mix10 (re-Conv^Mix10^) or with Mix4 (*A. caviae*, both *A. veronii* spp., *P. mossellii*) as a non-protective control (Suppl. Table S4). We tested genes encoding IL 1β (pro-inflammatory), IL22 (promoting gut repair), and IL10 (anti-inflammatory) cytokines. While we observed some variation in *il10* expression among non-infected re-conventionalized larvae, this did not correlate with protection. Furthermore, *il10* expression was not modulated by infection in any of the tested conditions (Fig. 4A). By contrast, we observed a strong induction of *il1b* and *il22* in GF zebrafish exposed to *F. columnare*^ALG^ (Fig. 4BC). However, although this induction was reduced in protected Conv, Re-Conv^Cm^ and Re-Conv^Mix10^, it was also observed in non-protective Re-Conv^Mix4^ larvae, indicating that down-modulation of the inflammatory response induced by *F. columnare* does not correlate with resistance to infection. Consistently, the use of a *myd88* mutant, a key adaptor of IL-1 and toll-like receptor signalling deficient in innate immunity(31, 46), showed that Conv or Re-Conv^Mix10^, but not GF *myd88* mutants survived *F. columnare* as well as wild-type zebrafish (Figure 4D). Moreover, *il1b* induction by *F. columnare* infection was observed only in GF larvae and was *myd88*-independent (Suppl. Figure S9). These results therefore indicated that the tested cytokine responses do not play a significant role in the microbiota-mediated protection against *F. columnare* infection.

**Figure 4.**
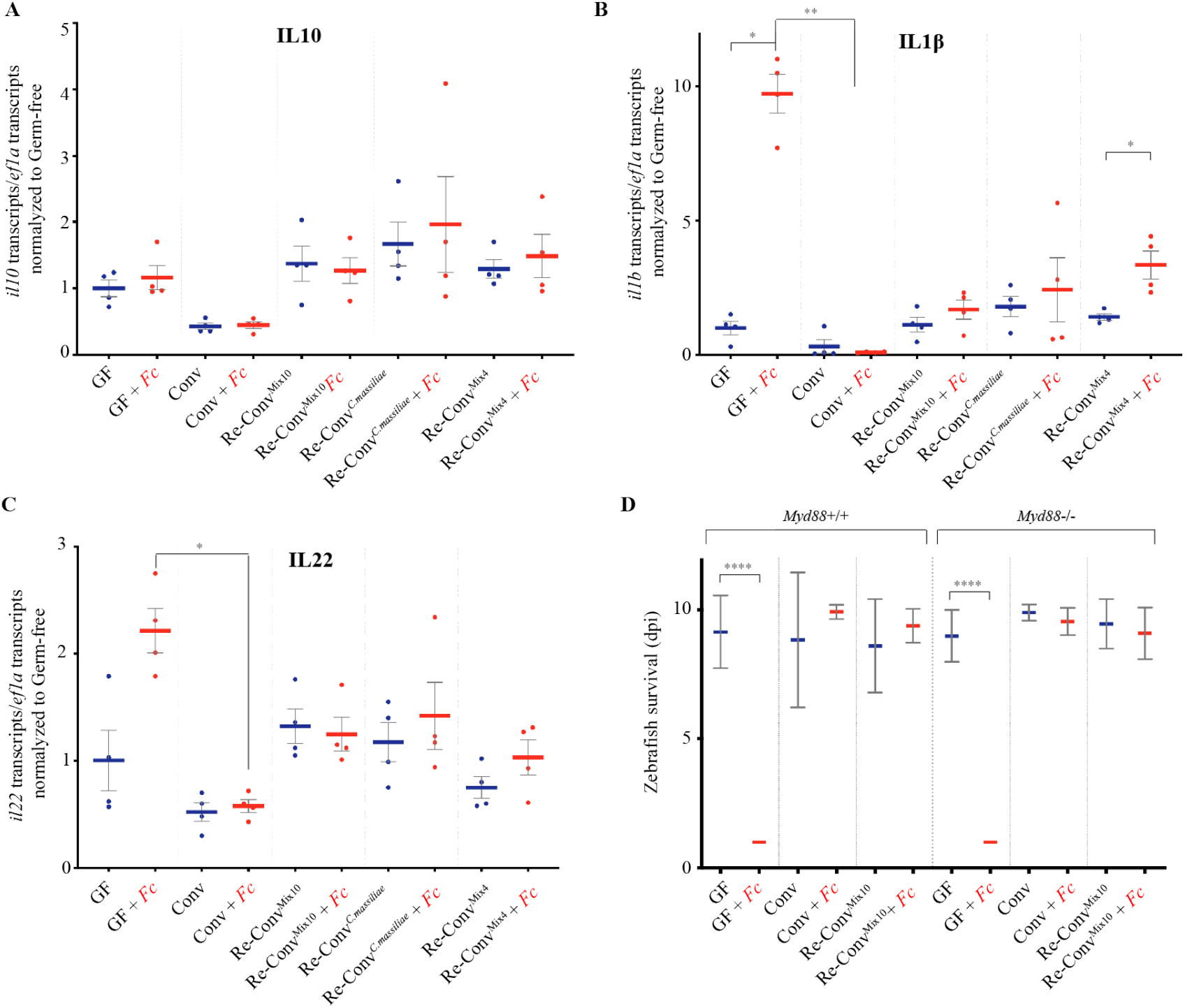
Zebrafish immune response to *F.columnare* infection. **A-C**: qRT-PCR analysis of host gene expression, 18 hours after exposure to *F. columnare,* in larvae re-conventionalized with indicated bacteria or bacterial mixes; each point corresponds to an individual larva. Expression of *il10* (**A**), *il1b* (**B**), and *il22* (**C**), by wild-type AB zebrafish; **D**: Comparison of the survival of *myd88−/−* and background-matched *myd88+/+* zebrafish after re-conventionalization and exposure to *F. columnare*^ALG^. Mean survival is represented by a thick horizontal bar with standard deviation. For each condition, n = 12 zebrafish larvae. Larvae mortality rate was monitored daily and surviving fish were euthanized at day 9 post exposition to the pathogen. A-D: Blue bars correspond to larvae not exposed to the pathogen and red mean bars correspond to exposed larvae. Indicated statistics correspond to unpaired, non-parametric Mann-Whitney test. ****: p<0.0001; **: p<0.005*: p<0.05, absence of *: non-significant.

### *C. massiliae* and Mix9 protect zebrafish from intestinal damages upon *F. columnare*^ALG^ infection

Histological analysis of GF larvae fixed 24h after exposure to *F. columnare*^ALG^ revealed extensive intestinal damage (Fig. 5A) prior to noticeable signs in other potential target organs such as gills or skin. To test the requirement for gut access in *F. columnare*^ALG^ infection process, we modified our standard rearing protocol of GF fish, which involves feeding with live germ-free *T. thermophila*. We found that, if left unfed, GF zebrafish did not die after *F. columnare*^ALG^ exposure, while feeding with either *T. thermophila* or another food source such as sterile fish food powder, restored sensitivity to *F. columnare*^ALG^ infection (Suppl. Fig S10), suggesting that successful infection requires feeding and ingestion.

**Figure 5.**
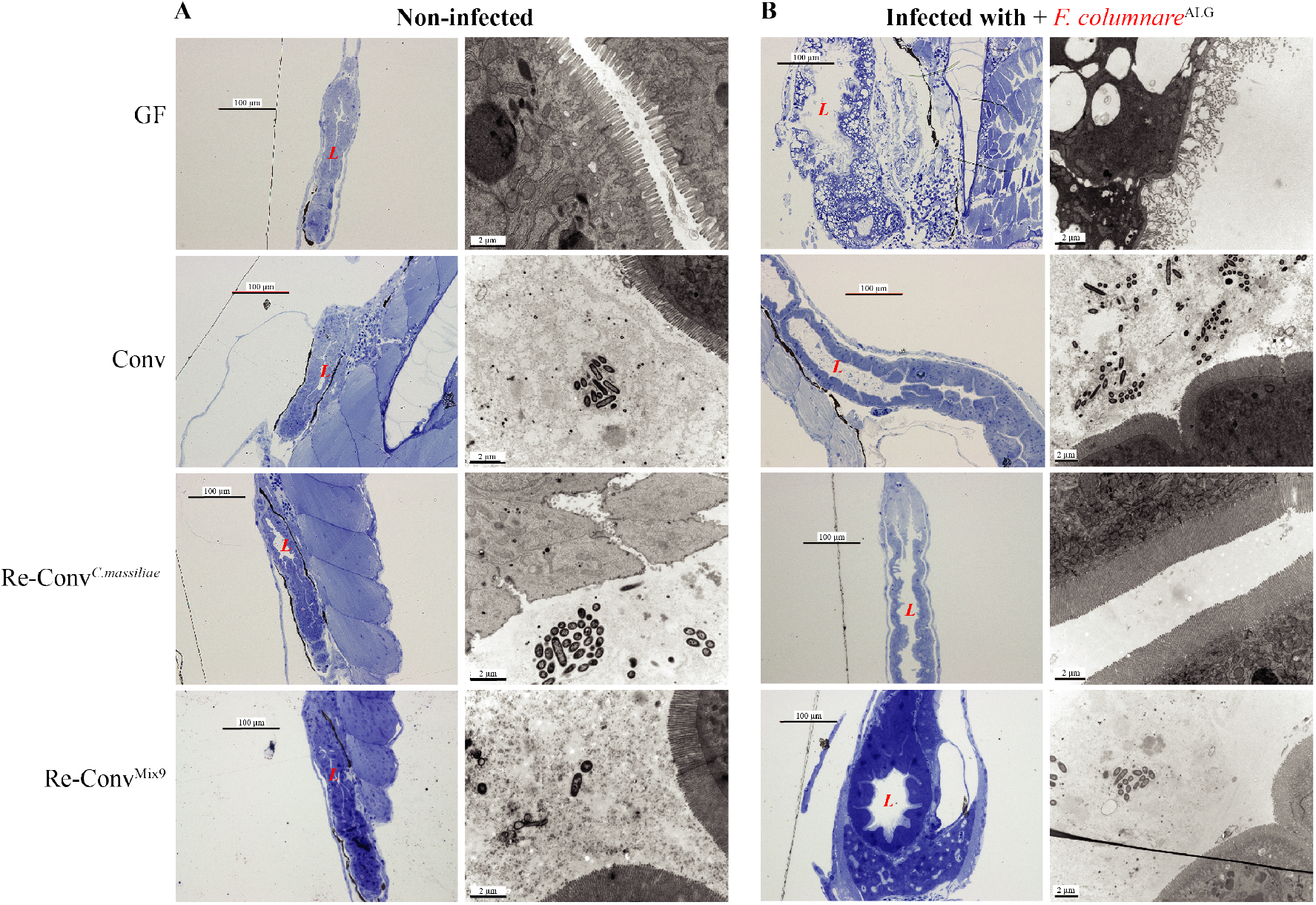
Intestine of infected germ-free zebrafish displays severe disorganization. Germ-free, conventional and re-conventionalized zebrafish larvae. Re-conventionalized zebrafish were inoculated at 4 dpf with Mix9 or *C. massiliae.* **A**: Representative picture of intestines of non-infected larvae. Fish were fixed for histology analysis or electron microscopy at 7 dpf. **B**: Representative picture of intestines of infected larvae exposed at 7 dpf to *F.columnare*^ALG^. In **A** and **B**: *Left column:* Toluidine blue staining of Epon-embedded zebrafish larvae for Light microscopy. *Right column*: Transmission electron microscopy. at 7 dpf (*right*). L= intestinal lumen.

Histological sections consistently showed severe disorganization of the intestine with blebbing in the microvilli and vacuole formation in *F. columnare*^ALG^-infected GF larvae (Fig. 5). In contrast, zebrafish pre-incubated with either *C. massiliae* or Mix9 consortium at 4 dpf, and then exposed to *F. columnare*^ALG^ at 6 dpf showed no difference compared to non-infected larvae or conventional infected larvae (Fig. 5), confirming full protection against *F. columnare*^ALG^ at the intestinal level.

### *C. massiliae* protects larvae and adult zebrafish against *F. columnare*

The clear protection provided by *C. massiliae* against *F. columnare*^ALG^ infection prompted us to test whether exogenous addition of this bacterium could improve microbiota-based protection towards this widespread fish pathogen. We first showed that zebrafish larvae colonized with *C. massiliae* were fully protected against all virulent *F. columnare* strains identified in this study (Fig. 6A). To test whether *C. massiliae* could also protect adult zebrafish from *F. columnare* infection, we pre-treated conventional 3-4-month-old Conv adult zebrafish with *C. massiliae* for 48h before challenging them with a high dose (5.10^6^ cfu/mL) *F. columnare*^ALG^. Monitoring of mortality rate showed that pre-treatment with *C. massiliae* significantly increased the survival rate of adult zebrafish upon *F. columnare*^ALG^ infection compared to non-treated conventional fish (p=0.0084 Mann-Whitney test, Figure 6B). Taken together, these results show that *C. massiliae* is a putative broad-spectrum probiotic protecting zebrafish against columnaris disease caused by *F. columnare*.

**Figure 6.**
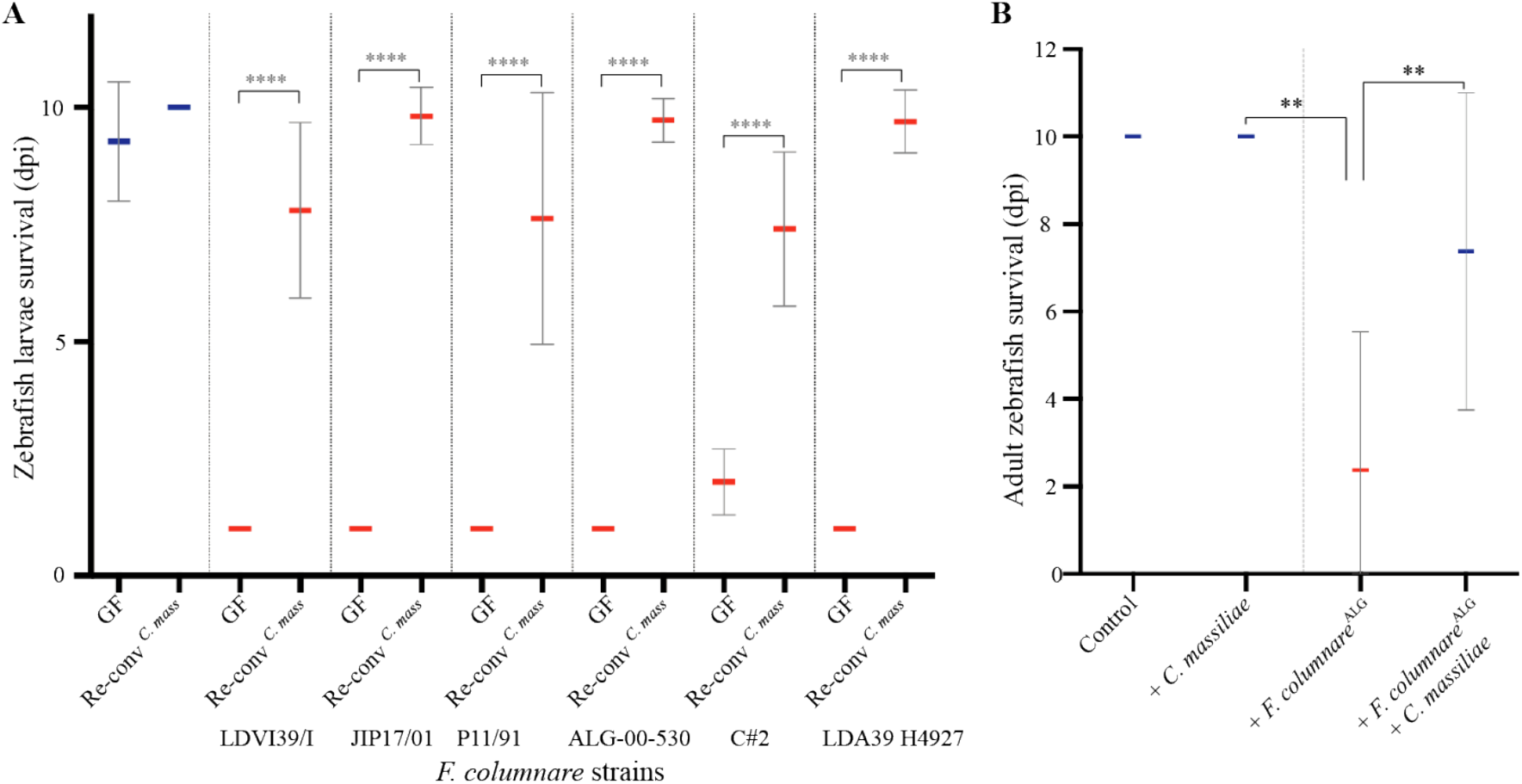
Pre-exposure to *C. massiliae* protects larval and adult zebrafish against *F. columnare* infection. **A:** Zebrafish larvae were inoculated at 4 dpf with 5.10^5^ cfu/mL of *C. massiliae* for 48h before infection at 6 dpf with virulent *F. columnare* strains. **B:** Survival of adult zebrafish with or without pre-exposure to *C. massiliae* (2.10^6^ cfu/mL for 48h) followed by exposure to *F. columnare*^ALG^(5.10^6^ cfu/mL for 1h) Mean survival is represented by a thick horizontal bar with standard deviation. For each condition, n = 12 zebrafish larvae or 10 adult. Zebrafish mortality rate was monitored daily and surviving fish were euthanized at day 9 post exposition to the pathogens. Blue bars correspond to larvae not exposed to the pathogen and red mean bars correspond to exposed larvae. Indicated statistics correspond to unpaired, non-parametric Mann-Whitney test. ****: p<0.0001; **: p<0.005; absence of *: non-significant.

## DISCUSSION

In this study, we used re-conventionalization of otherwise germ-free zebrafish larvae to show that conventional-level protection against infection by a broad range of highly virulent *F. columnare* strains is provided by a set of 10 culturable bacterial strains, belonging to 9 different species, isolated from the indigenous standard laboratory zebrafish microbiota. With the exception of the Bacteroidetes *C. massiliae*, this protective consortium was dominated by Proteobacteria such as *Pseudomonas* and *Aeromonas* spp., bacteria commonly found in aquatic environments (47, 48). Despite the relative permissiveness of zebrafish larvae microbiota to environmental variations and inherent variability between samples (49), we showed that these ten bacteria were consistently identified in four different zebrafish facilities, suggesting the existence of a core microbiota with important colonization resistance functionality. Use of controlled combinations of these 10 bacterial species enabled us to show a very robust species-specific protection effect in larvae mono-associated with *C. massiliae.* We also identified a community-level protection provided by the combination of the 9 other species that were otherwise unable to protect against *F. columnare* when provided individually. This protection was however less reproducible and required a minimum inoculum of 5.10^4^ cfu/mL, compared to 5.10^2^ cfu/mL with *C. massiliae*. These results therefore suggest the existence of two distinct microbiota-based protection scenarios: one based on a membership effect provided by *C. massiliae*, and the other mediated by the higher-order activity of the Mix9 bacterial community.

Although protection against *F. columnare* infection does not seem to rely on microbiota-based immuno-modulation, we cannot exclude that, individually, some members of the studied core zebrafish microbiota could induce pro- or anti-inflammatory responses masked in presence of the full Mix10 consortium (1). We also acknowledge that the mechanistic aspects of the two identified colonization resistance scenarios are still unclear and further studies are required to determine whether they involve nutrient depletion, adhesion inhibition, direct or indirect growth inhibition leading to niche exclusion or any other process (6, 13). In the case of the protection provided by *C. massiliae* against *F. columnare,* these two phylogenetically close bacteria could compete for similar resources and directly antagonize each other, as shown to occur between *Bacteroidetes* species (50–53). Interestingly, infected larvae re-conventionalized with either C. *massiliae* or Mix9 showed no signs of the intestinal damage displayed by germ-free larvae, suggesting that both *C. massiliae* and Mix9 provide similar intestinal resistance to *F. columnare* infection. Whereas microbial colonization contributes to gut maturation and stimulates the production of epithelial passive defences such as mucus (54, 55), lack of intestinal maturation is unlikely to be contributing to *F. columnare*-induced mortality, as mono-colonized larvae or larvae re-conventionalized with non-protective mixes died as rapidly as germ-free larvae.

Several studies have monitored the long-term assembly and development of the zebrafish microbiota from larvae to sexually mature adults, however little is known about the initial colonization establishment of the larvae after hatching (56, 57). Neutral (stochastic) and deterministic (host niche-based) processes (58–60) lead to microbial communities that are mostly represented by a limited number of highly abundant species with highly diverse low-abundant populations. In our experiments, the Mix10 species inoculum corresponded to an equiratio bacterial mix, thus starting from an engineered and assumed total evenness (E = 1) (61, 62). Evenness was still relatively high (0.84) and remained similar up until 8h in our study, indicating that most of the ten species were able to colonize the larvae. From the perspective of community composition, a loss of diversity is often associated with decreased colonization resistance, but it remains unclear whether this increased susceptibility is due to the loss of certain key member species of the microbial community and/or a change in their prevalence (3, 9).

We further investigated resistance to infection by exposing established bacterial communities to different antibiotic perturbations, followed by direct challenge with *F. columnare* (to study core microbiota sensitivity to disturbance) or after recovery (to study its resilience) (12, 63). Antibiotics are known to shift the composition and relative abundances of the microbiota according to their spectrum (13, 64). We observed that penicillin/streptomycin treatment that would affect most of the core species, reduced the abundance of all but two species (*A. veronii* 1 and *P. myrsinacearum*) that became relatively dominant during recovery, but failed to provide protection against *F. columnare*. With the kanamycin treatment, colonization resistance was fully restored at the end of the 24h recovery period, indicative of a resilience that could result from species recovering quickly to their pre-perturbation levels due to fast growth rates, physiological flexibility or mutations (65). Interestingly, even taking into account potential biases associated with the use of the 16S rRNA as a proxy index to determine relative abundance (66, 67), evenness was similarly reduced during recovery for both treatments, but abundance at phylum level changed to 48% for Proteobacteria, and 52% for Bacteroidetes compared to the >98% of Proteobacteria with the penicillin/streptomycin treatment. Furthermore, *C. massiliae* was detected as rare (<1%) in conventional larvae, suggesting that it could have a disproportionate effect on the community or that community-level protection provided by the nine other bacteria was also responsible for the protection of conventional larvae to *F. columnare* infection.

We showed that germ-free zebrafish larvae are highly susceptible to a variety of different *F. columnare* genomovars isolated from different hosts, demonstrating that they are a robust animal model for the study of its pathogenicity. Recently, *F. columnare* mutants in Type 9 secretion system (T9SS) were shown to be avirulent in adult zebrafish, suggesting that proteins secreted by the T9SS are likely to be key, but still largely unidentified, *F. columnare* virulence determinants (45). Body skin, gills, fins and tail are also frequently damaged in salmonid fish, whereas severe infection cases are associated with septicemia (68). We could not identify such clear *F. columnare* infection sites in zebrafish larvae, perhaps due to the very low dose of infection used, with less than 100 cfu recovered from infected moribund larvae. However, several lines of evidence suggest that the gut is the main target of *F. columnare* infection in our model: (i) unfed germ-free larvae survived exposure, (ii) histology analysis showing severe disruption of the intestinal region just hours after infection in germ-free larvae, and (iii) induction of *il22* in germ-free larvae exposed to *F. columnare*, since a major function of IL-22 is to promote gut repair (69). This induction appears to be a consequence of the pathogen-mediated damage, as there was no observed induction in conventional or re-conventionalized larvae. The very rapid death of larvae likely caused by this severe intestinal damage may explain why other common target organs of columnaris disease showed little damage.

In this study, we showed that *C. massiliae* is a promising probiotic candidate that could contribute to reduce the use of antibiotic to prevent columnaris diseases in research and aquaculture settings. Whereas *C. massiliae* provided full and robust protection against all tested virulent *F. columnare* genomovars and significantly increased survival of exposed adult conventional zebrafish, further studies are needed to elucidate *C. massiliae* protection potential in other teleost fish. However, the endogenous nature of *C. massiliae* suggests that it could establish itself as a long-term resident of the zebrafish larval and adult microbiota, an advantageous trait when seeking a stable modulation of the bacterial community over long periods (33).

In conclusion, the use of a simple and tractable zebrafish larval model to mine indigenous host microbial communities allowed us to identify two independent protection pathways against the same pathogen. Whereas further study will determine how these pathways may contribute to protection against a wider range of pathogens, this work also provides insights into how to the engineering of stable protective microbial communities with controlled colonization resistance functions.

## ACKNOWLEDGEMENTS

We thank Mark McBride, Pierre Boudinot and Rebecca Stevick for critical reading of the manuscript. We are grateful to the late Covadonga Arias for the gift of *F. columnare* ALG 00-530, to Mark McBride for *F. columnare* C#2 strain and to Jean-François Bernardet for all other *F. columnare* strains. Prof. Annemarie Meijer (Leiden University) kindly provided the *myd88* mutant zebrafish line. We thank Julien Burlaud-Gaillard and Rustem Uzbekov the IBiSA Microscopy facility, Tours University, France and the following zebrafish facility teams for providing eggs: José Perez and Yohann Rolin(Institut Pasteur), Nadia Soussi-Yanicostas (INSERM Robert Debré), Sylvie Schneider-Manoury and Isabelle Anselme (UMR7622, University Paris 6) and Frédéric Sohm (AMAGEN Gif-sur-Yvette).

## FUNDING

This work was supported by the Institut Pasteur, the French Government’s *Investissement d’Avenir* program: *Laboratoire d’Excellence* ‘Integrative Biology of Emerging Infectious Diseases’ (grant no. ANR-10-LABX-62-IBEID to J.M.G.), the Fondation pour la Recherche Médicale (grant no. DEQ20180339185 to J.M.G.). F.S. was the recipient of a post-doctoral Marie Curie fellowship from the EU-FP7 program, J.B.B. was the recipient of a long-term post-doctoral fellowship from the Federation of European Biochemical Societies (FEBS) and by the European Union’s Horizon 2020 research and innovation programme under the Marie Skłodowska-Curie grant agreement No 842629. D.P-P was supported by an Institut Carnot MS Postdoctoral fellowship.

The funders had no role in study design, data collection and analysis, decision to publish, or preparation of the manuscript.

## CONFLICT OF INTEREST

The authors of this manuscript have the following conflict of interest: a provisional patent application has been filed: “*bacterial strains for use as probiotics, compositions thereof, deposited strains and method to identify probiotic bacterial strains*” by J.-M.G, F. A. S., D.P.-P. and J. B. B. The other authors declare no conflict of interest in relation to the submitted work.

## AUTHOR CONTRIBUTIONS

F. A. S., J. B. B., D.P.-P., J.-P. L. and J.-M.G. designed the experiments. O.R. contributed to the initial experiments. V.B. and J.-P. L. provided zebrafish material and advice. F. A. S, J. B. B., D.P.-P., B.A., V.B. and J.-P. L. performed the experiments. S.B., S.H. performed bacterial genome sequencing and analysis, A.G., S.V., F. A. S and D.P.-P. performed the bioinformatic and sequence analyses. F. A. S, J. B. B., D.P.-P., J.-P. L. and J.-M.G. analysed the data and wrote the paper with significant contribution from O.R. and E.D..

