## supplementary figures S1-to S10 and Supplementary Tables S1 to S4 for "Mining zebrafish microbiota reveals key community-level resistance against fish pathogen infection"

### SUPPLEMENTARY INFORMATION

#### SUPPLEMENTARY FIGURES (supplementary figures 1-10.)

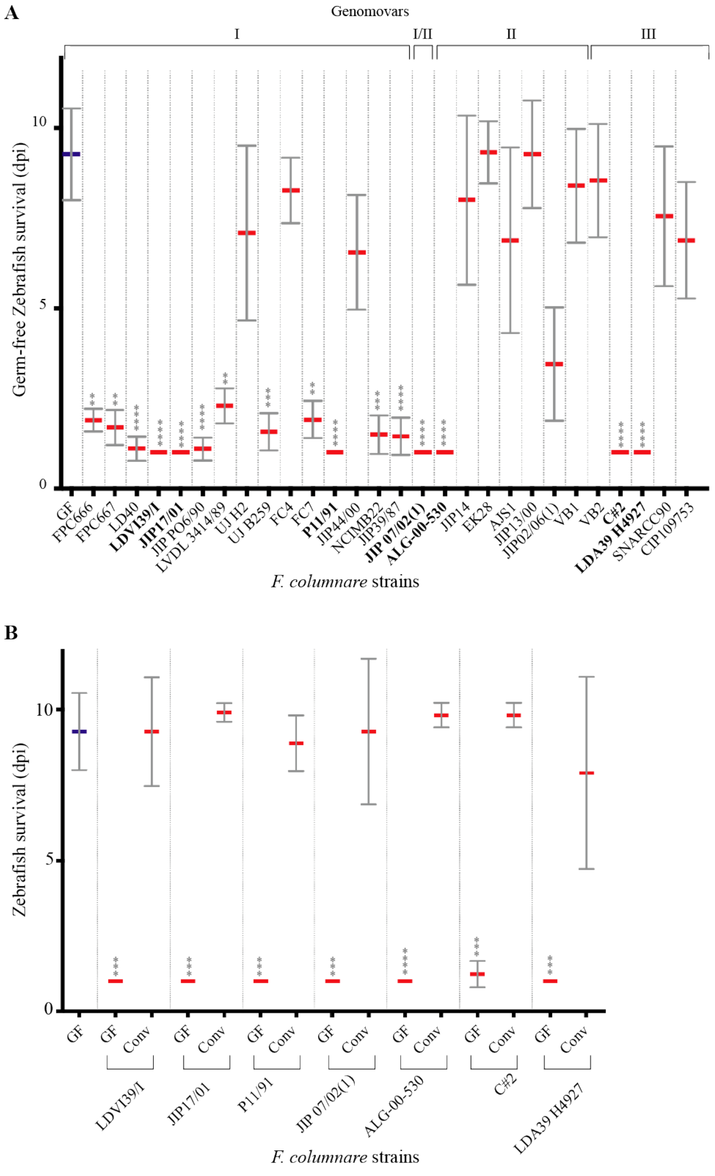


**Supplementary Figure S1**. **Screen for *F. columnare* strains killing germ-free zebrafish**. **A**. Survival of GF zebrafish larvae exposed to a collection of 28 strains of *F. columnare*. **B**. Survival of GF and Conv zebrafish larvae exposed to the 7 most virulent *F. columnare* strains. Zebrafish larvae were infected at 6 dpf (= 0 dpi) by immersion for 3h with $5.10^5$ cfu/mL. Mean survival is represented by a thick horizontal bar with standard deviation. For each condition, n = 12 zebrafish larvae. Larvae mortality rate was monitored daily and surviving fish were euthanized at day 9 post infection. Statistics correspond to unpaired, non-parametric Mann-Whitney test comparing all conditions to non-infected GF. ****: p<0.0001; ***: p<0.001; absence of star: non-significant. Blue mean bars correspond to larvae not exposed to the pathogen and red mean bars correspond to exposed larvae.

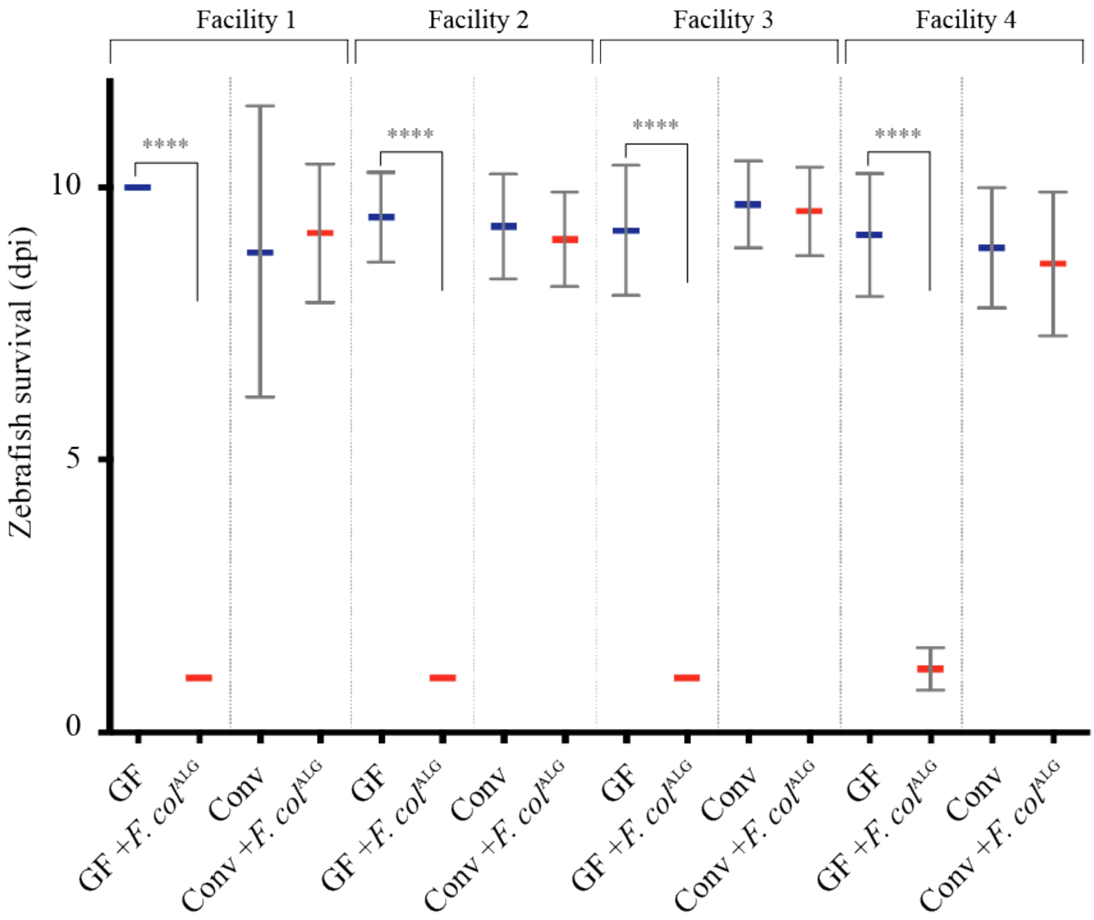


**Supplementary Figure S2. Conventional larvae from 4 different zebrafish facilities are also protected against *F. columnare* infection.** Zebrafish AB line eggs from 4 different zebrafish facilities were collected: Facility 1 – Hospital Robert Debré academic facility, Paris, France; Facility 2 and 3 – two academic facilities at University Paris 6, Paris, France; Facility 4 – commercial Amagen facility in Gif-sur-Yvette, France. 6 dpf (= 0 dpi) GF or Conv zebrafish larvae from each facility were exposed to *F. columnare*$^{ALG}$ by bath immersion and transferred after 3 hours to sterile water. Mean survival is represented by a thick horizontal bar with standard deviation. For each condition, n = 12 zebrafish larvae. Blue mean bars correspond to larvae not exposed to the pathogen and red mean bars correspond to exposed larvae. Larvae mortality rate was monitored daily and surviving fish were euthanized at day 10 post infection. Indicated statistics correspond to unpaired, non-parametric Mann-Whitney test. ****: $p<0.0001$; absence of *: non-significant.

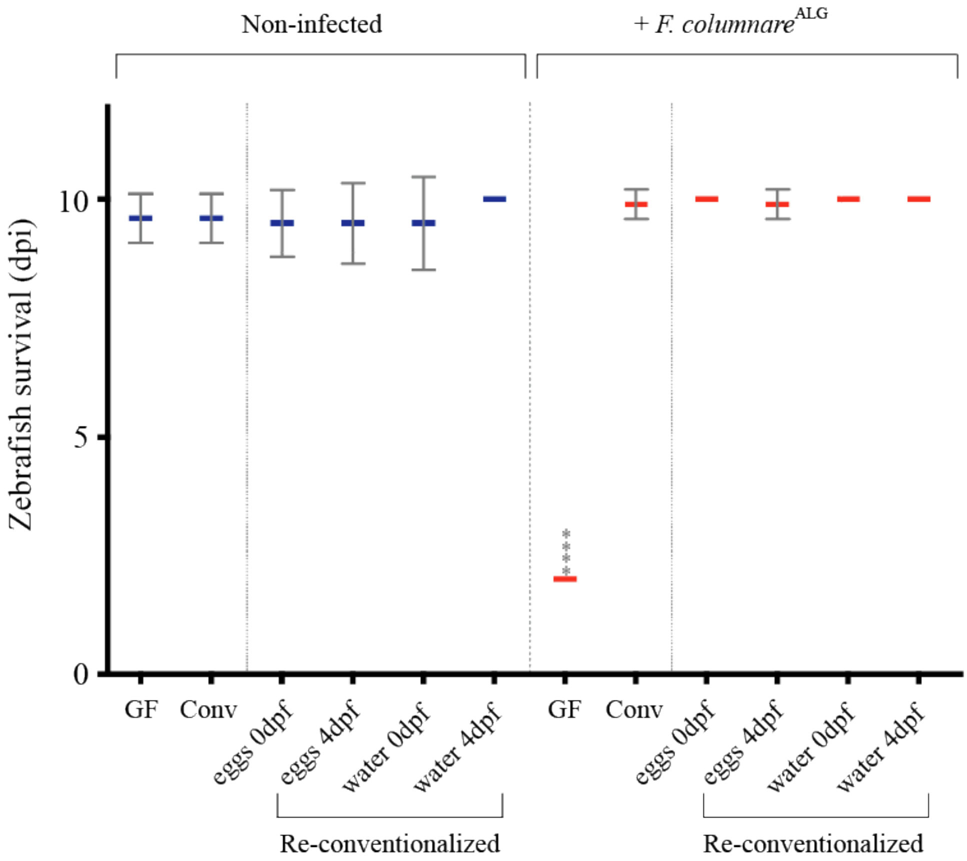


**Supplementary Figure S3. Reconventionalization of germ-free zebrafish larvae confers protection against *F. columnare*.**

Resistance to *F. columnare*$^{ALG}$ of GF and Conv zebrafish larvae placed in contact with fish facility tank water or mashed non-sterile eggs at 0 (sterilization day) or 4 dpf (hatching day). *F. columnare*$^{ALG}$ inoculum doses = $5.10^5$ cfu/mL. Mean survival is represented by a thick horizontal bar with standard deviation. For each condition, n = 12 zebrafish larvae. Larvae mortality rate was monitored daily and surviving fish were euthanized at day 10 post exposition to the pathogen. Statistics correspond to unpaired, non-parametric Mann-Whitney test comparing all conditions to non-exposed GF. ****: $p<0.0001$; absence of *: non-significant. Blue mean bars correspond to larvae not exposed to the pathogen and red mean bars correspond to exposed larvae.

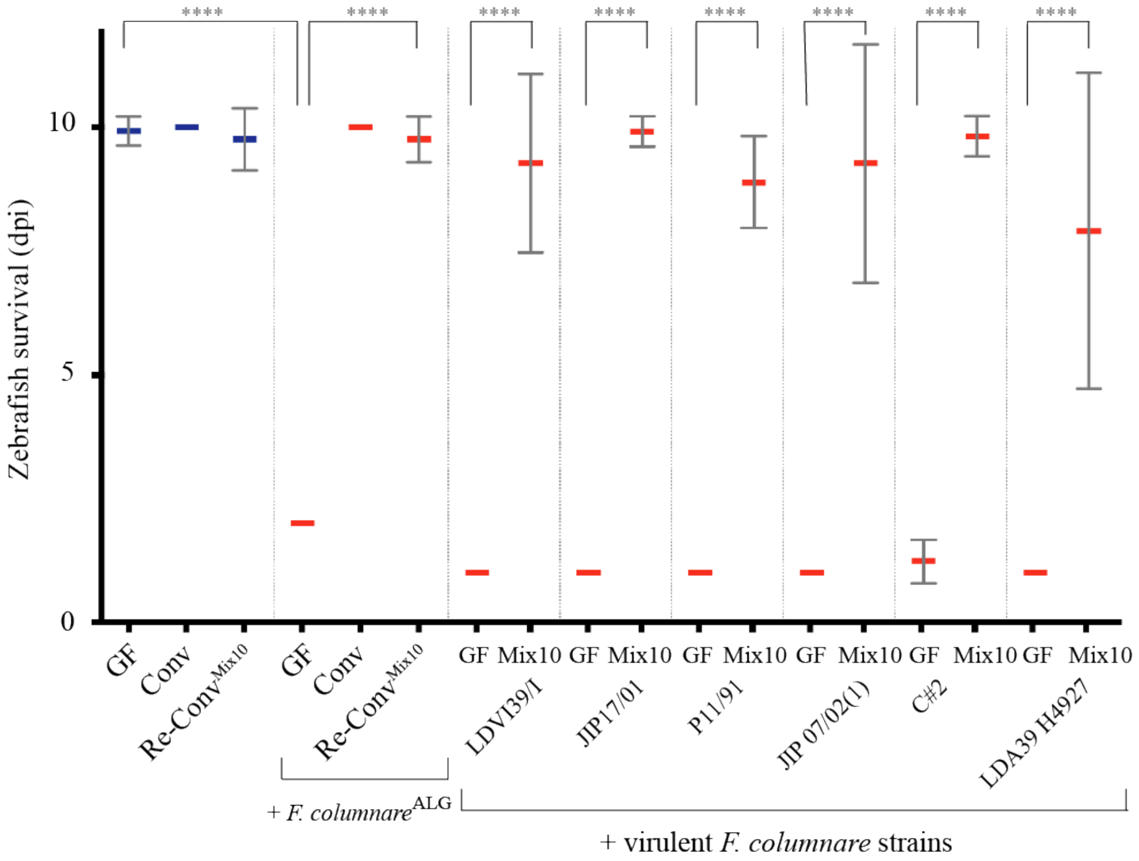


**Supplementary Figure S4. A mix of the 10 culturable bacteria constituting the core zebrafish larvae microbiota confers protection against *F. columnare*.** The 10 culturable strains identified as the core conventional microbiota were added at an equivalent concentration of $5.10^5$ cfu/mL mix to 4 dpf larvae followed by infection challenge at 6 dpf. Mean survival is represented by a thick horizontal bar with standard deviation. For each condition, n = 12 zebrafish larvae. Larvae mortality rate was monitored daily and surviving fish were euthanized at day 10 post exposition to the pathogen. Indicated statistics correspond to unpaired, non-parametric Mann-Whitney test. ****: p<0.0001; absence of *: non-significant. Blue mean bars correspond to larvae not exposed to the pathogen and red mean bars correspond to exposed larvae.

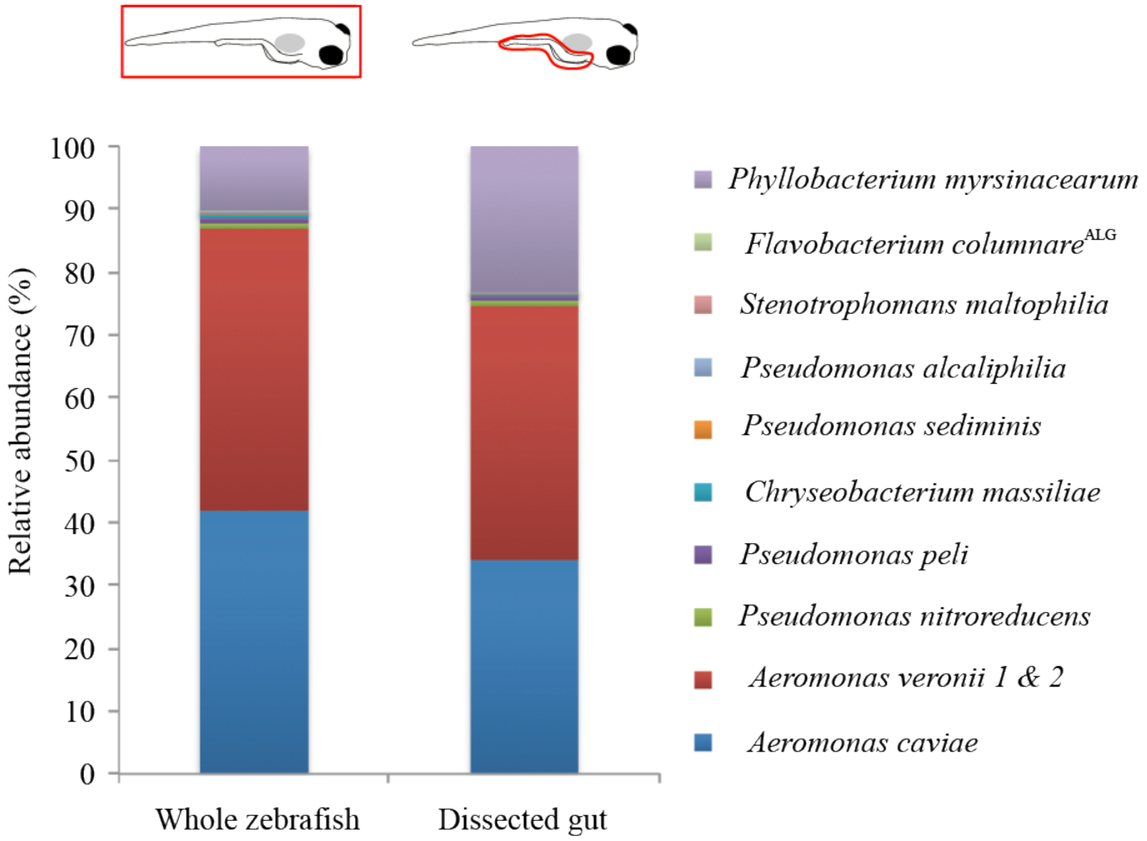


**Supplementary Figure S5. Illumina sequencing comparison of gut and whole larval intestinal bacterial content.** Larvae re-conentionalized with Mix10 and infected with *F. columnare*$^{ALG}$ at 6 dpf for 3h were euthanized and washed. DNA was extracted from pools of 10 whole larvae or of pools of 10 intestinal tubes dissected with sterile surgical tweezer and subjected to Illumina 16S rRNA gene sequencing. GF larvae and dissected GF intestines were sampled as controls. No statistically significant difference was found between whole fish and gut bacterial content ($p$ = 0.99, two-tailed t-test of OTU abundances). Entire larvae were therefore used in the experiment monitoring bacterial establishment and recovery.

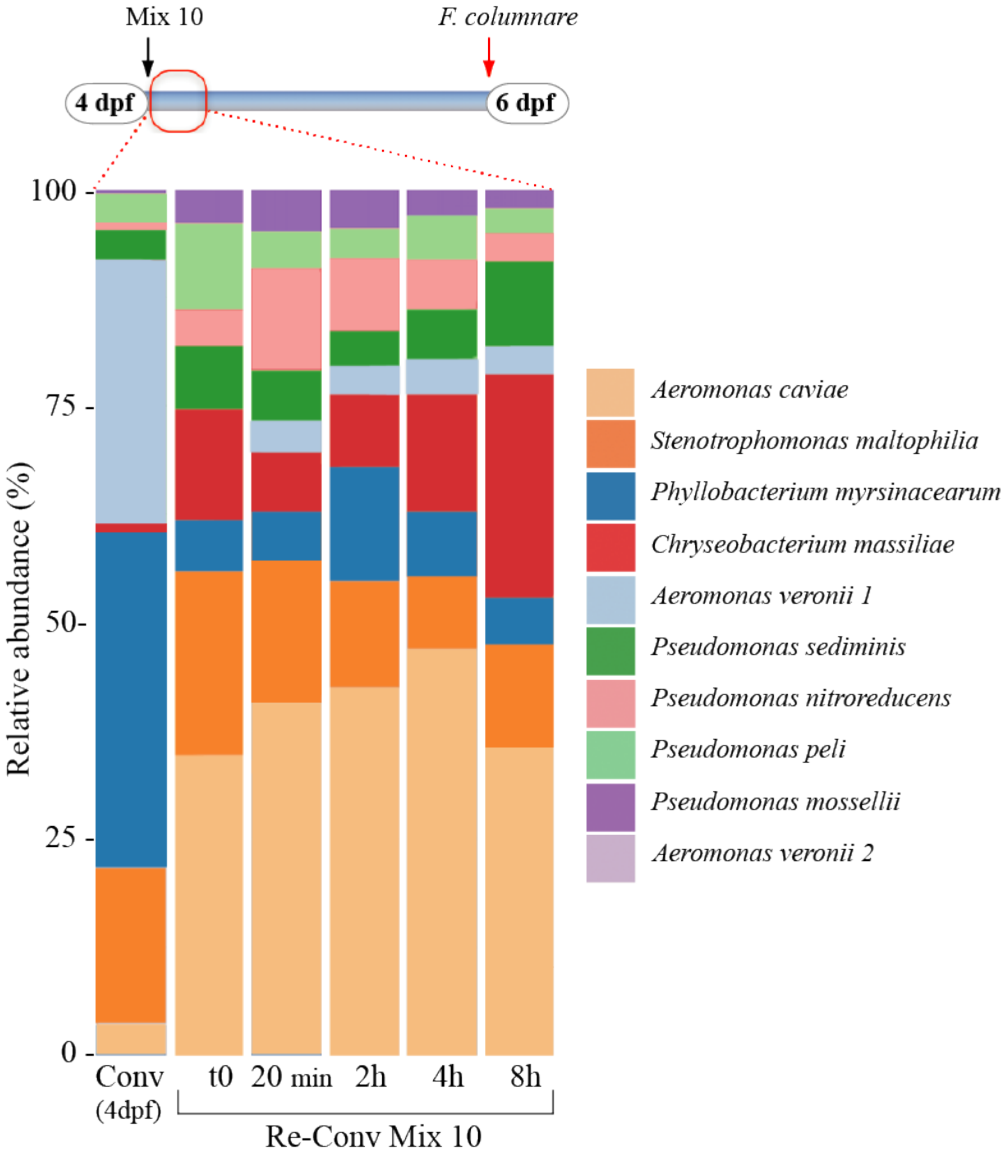


**Supplementary Figure S6. Dynamics of bacterial community establishment in reconventionalized zebrafish larvae.** Relative abundances of the 10 species composing the identified Mix10 core protective zebrafish microbiota at different time points using 16S rRNA gene amplicon sequencing of pools of 10 larvae. Left bar represents the relative abundance of microbiota species of conventional zebrafish larvae on hatching day (4 dpf). For Re-Conv Mix10 populations, *Tetrahymena*-fed GF larvae were incubated with an equiratio combination of the 10 species ($5.10^5$ cfu/mL each) composing the microbiota core at 4 dpf.

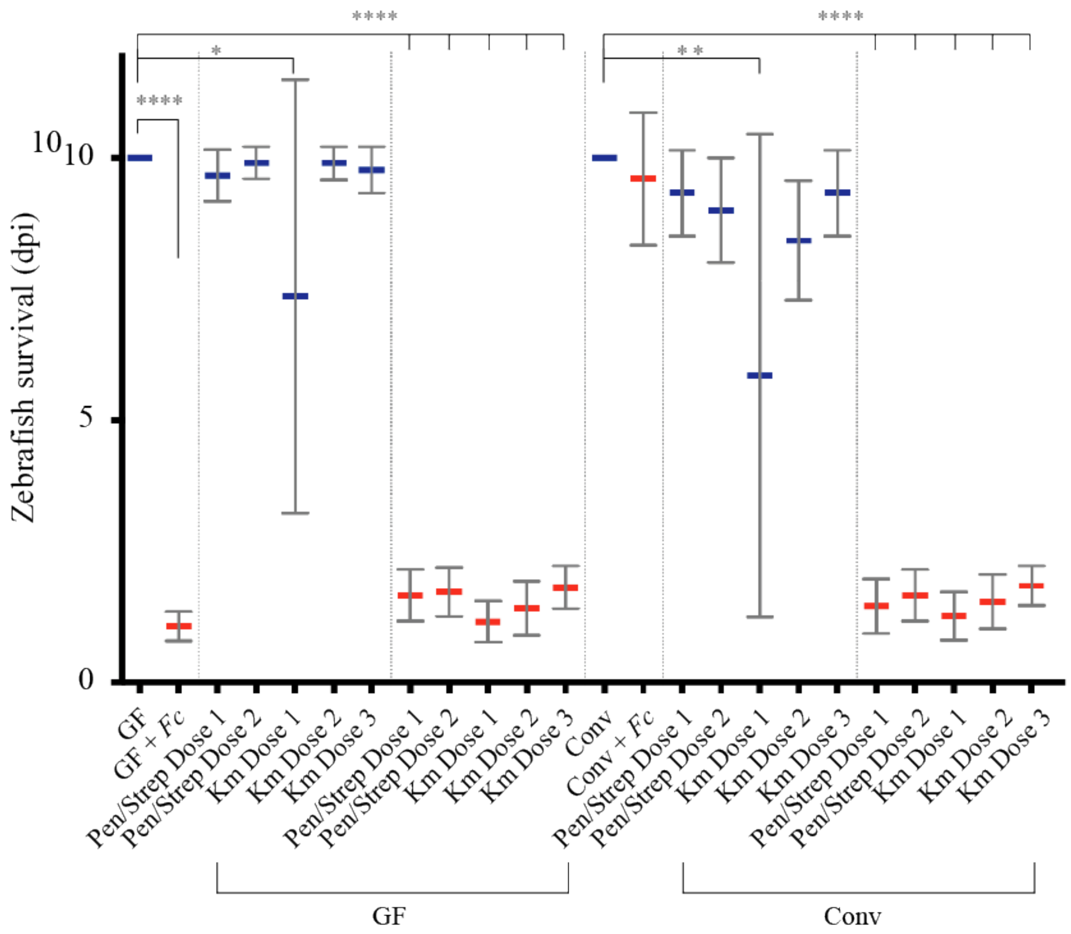


**Supplementary Figure S7. Effect of antibiotic treatment on zebrafish survival.** GF and Conv zebrafish larvae were exposed to antibiotic treatment for 16 hours at 4 dpf. Antibiotics were then washed off and the zebrafish larvae were infected with *F. columnare*$^{ALG}$. Different concentrations of penicillin/streptomycin or kanamycin were used to identify a non-toxic antibiotic treatment causing microbiota dysbiosis. Penicillin/streptomycin dose 1= 250µg/mL; dose 2= 15.6 µg/mL; kanamycin dose 1= 200 µg/mL; dose 2= 50 µg/mL; dose 3= 25 µg/mL. Mean survival is represented by a thick horizontal bar with standard deviation. For each condition, n = 12 zebrafish larvae. Blue mean bars correspond to larvae not exposed to the pathogen and red mean bars correspond to exposed larvae. Larvae mortality rate was monitored daily and surviving fish were euthanized at day 10 post infection. Indicated statistics correspond to unpaired, non-parametric Mann-Whitney test. ****: p<0.0001; **: p<0.005; *: p<0.05, absence of *: non-significant.

A

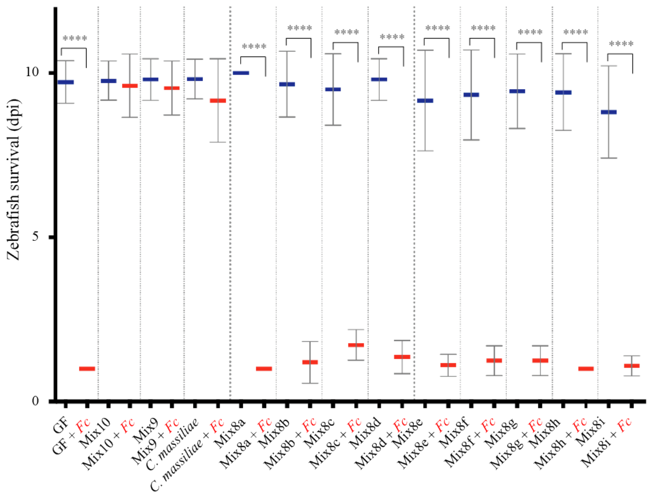


B

| Mix8 combinations | | | | |
|---|---|---|---|---|
| **Mix8a** | **Mix8b** | **Mix8c** | **Mix8d** | **Mix8e** |
| *A. veronii* 1 | *A. veronii* 1 | *A. veronii* 1 | *A. veronii* 1 | *A. veronii* 1 |
| *P. mossellii* | *P. mossellii* | *P. mossellii* | *P. mossellii* | *P. mossellii* |
| *S. maltophilia* | *S. maltophilia* | *S. maltophilia* | *S. maltophilia* | *S. maltophilia* |
| *A. caviae* | *A. caviae* | *A. caviae* | *A. caviae* | - |
| *P. peli* | *P. peli* | *P. peli* | - | *P. peli* |
| *P. sediminis* | *P. sediminis* | - | *P. sediminis* | *P. sediminis* |
| *P. myrsinacearum* | - | *P. myrsinacearum* | *P. myrsinacearum* | *P. myrsinacearum* |
| - | *A. veronii* 2 | *A. veronii* 2 | *A. veronii* 2 | *A. veronii* 2 |
| *P. nitroreducens* | *P. nitroreducens* | *P. nitroreducens* | *P. nitroreducens* | *P. nitroreducens* |
| | **Mix8f** | **Mix8g** | **Mix8h** | **Mix8i** |
| | *A. veronii* 1 | *A. veronii* 1 | - | *A. veronii* 1 |
| | *P. mossellii* | - | *P. mossellii* | *P. mossellii* |
| | - | *S. maltophilia* | *S. maltophilia* | *S. maltophilia* |
| | *A. caviae* | *A. caviae* | *A. caviae* | *A. caviae* |
| | *P. peli* | *P. peli* | *P. peli* | *P. peli* |
| | *P. sediminis* | *P. sediminis* | *P. sediminis* | *P. sediminis* |
| | *P. myrsinacearum* | *P. myrsinacearum* | *P. myrsinacearum* | *P. myrsinacearum* |
| | *A. veronii* 2 | *A. veronii* 2 | *A. veronii* 2 | *A. veronii* 2 |
| | *P. nitroreducens* | *P. nitroreducens* | *P. nitroreducens* | - |

C

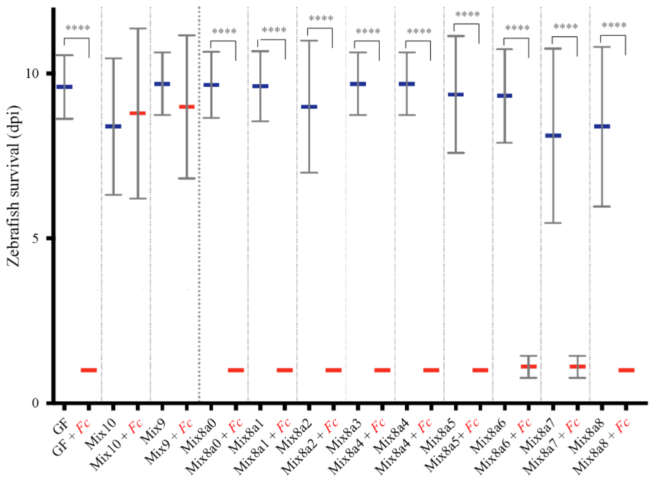


D

| Mix8a combinations | | | | |
|---|---|---|---|---|
| **Mix8a-0** | **Mix8a-1** | **Mix8a-2** | **Mix8a-3** | **Mix8a-4** |
| - | ***A. veronii* 1** | ***P. mossellii*** | ***S. maltophilia*** | ***A. caviae*** |
| *A. veronii* 1 | ***A. veronii* 1** | *A. veronii* 1 | *A. veronii* 1 | *A. veronii* 1 |
| *P. mossellii* | *P. mossellii* | ***P. mossellii*** | *P. mossellii* | *P. mossellii* |
| *S. maltophilia* | *S. maltophilia* | *S. maltophilia* | ***S. maltophilia*** | *S. maltophilia* |
| *A. caviae* | *A. caviae* | *A. caviae* | *A. caviae* | ***A. caviae*** |
| *P. peli* | *P. peli* | *P. peli* | *P. peli* | *P. peli* |
| *P. sediminis* | *P. sediminis* | *P. sediminis* | *P. sediminis* | *P. sediminis* |
| *P. myrsinacearum* | *P. myrsinacearum* | *P. myrsinacearum* | *P. myrsinacearum* | *P. myrsinacearum* |
| *P. nitroreducens* | *P. nitroreducens* | *P. nitroreducens* | *P. nitroreducens* | *P. nitroreducens* |
| | **Mix8a-5** | **Mix8a-6** | **Mix8a-7** | **Mix8a-8** |
| | ***P. peli*** | ***P. sediminis*** | ***P. myrsinacearum*** | ***P. nitroreducens*** |
| | *A. veronii* 1 | *A. veronii* 1 | *A. veronii* 1 | *A. veronii* 1 |
| | *P. mossellii* | *P. mossellii* | *P. mossellii* | *P. mossellii* |
| | *S. maltophilia* | *S. maltophilia* | *S. maltophilia* | *S. maltophilia* |
| | *A. caviae* | *A. caviae* | *A. caviae* | *A. caviae* |
| | ***P. peli*** | *P. peli* | *P. peli* | *P. peli* |
| | *P. sediminis* | ***P. sediminis*** | *P. sediminis* | *P. sediminis* |
| | *P. myrsinacearum* | *P. myrsinacearum* | ***P. myrsinacearum*** | *P. myrsinacearum* |
| | *P. nitroreducens* | *P. nitroreducens* | *P. nitroreducens* | ***P. nitroreducens*** |

**Supplementary Figure S8. Combinations of 8 species from the protective Mix9 do not protect against *F. columnare* infections. A:** Zebrafish larvae survival to *F. columnare*$^{ALG}$ infection at 6 dpf after reconventionalization with different possible mixes including 8 species of the Mix9 consortium. Mix10= Re-ConvMix$^{10}$, Mix9= Re-Conv$^{Mix9}$, Mix8=Re-Conv$^{Mix8}$. **B:** Table showing different combinations used for the reconventionalization at 4 dpf. **C**: Zebrafish larvae survival to *F. columnare*$^{ALG}$ infection at 6 dpf after reconventionalization with eight different combinations of the non-protective Mix8a were tested, in which the quantity of 1 of the 8 species was doubled in each combination (indicated in yellow in **D**). **D**: Table showing different combinations used for the reconventionalization at 4 dpf. Mix10= Re-ConvMix$^{10}$, Mix9= Re-Conv$^{Mix9}$, Mix8= Re-Conv$^{Mix8}$. **A** and **C**: Mean survival is represented by a thick horizontal bar with standard deviation. For each condition, n = 12 zebrafish larvae. Blue mean bars correspond to larvae not exposed to the pathogen and red mean bars correspond to exposed larvae. Larvae mortality rate was monitored daily and surviving fish were euthanized at day 10 post infection. Indicated statistics correspond to unpaired, non-parametric Mann-Whitney test. ****: p<0.0001; absence of *: non-significant.

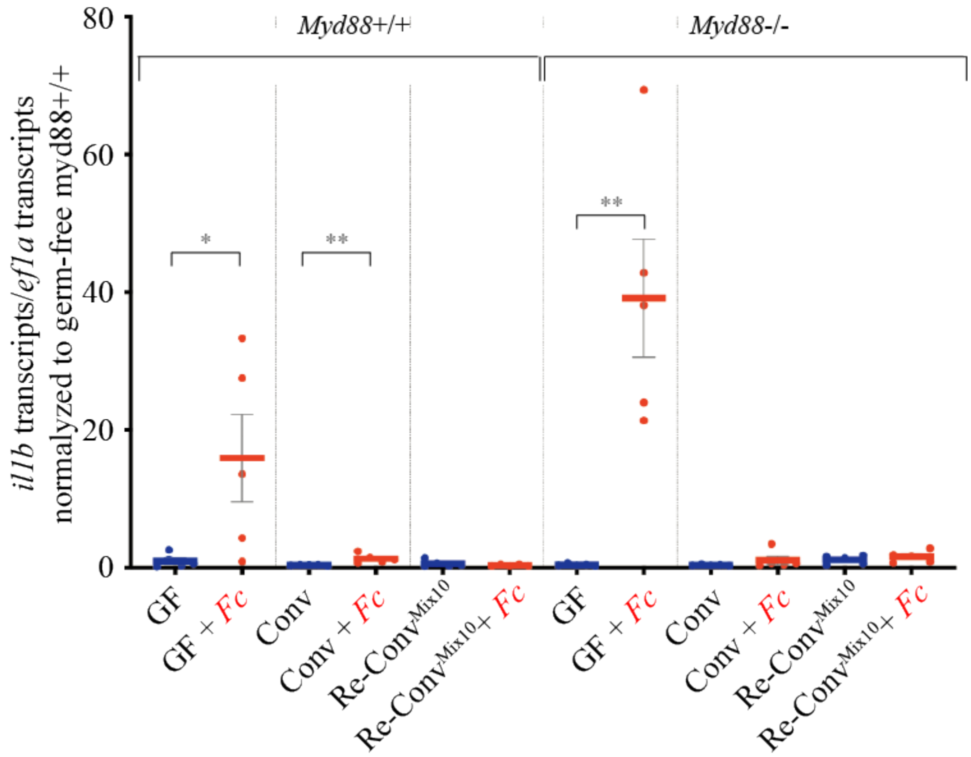


**Supplementary Figure S9. Expression of *il1b* in WT and myd88$^{-/-}$ zebrafish mutants reconventionalized with indicated various bacteria or bacterial mixes.** qRT-PCR analysis of *il1b* gene expression were performed 18 hours after exposure to *F.columnare*$^{ALG}$. Each point corresponds to an individual larva. Blue mean bars correspond to larvae not exposed to the pathogen and red mean bars correspond to exposed larvae. Larvae mortality rate was monitored daily and surviving fish were euthanized at day 10 post infection. Indicated statistics correspond to unpaired, non-parametric Mann-Whitney test. *: $p<0.005$*: $p<0.05$; absence of *: non-significant. Mean survival is represented by a thick horizontal bar. Blue bars correspond to non-infected larvae and red bars correspond to infected zebrafish.

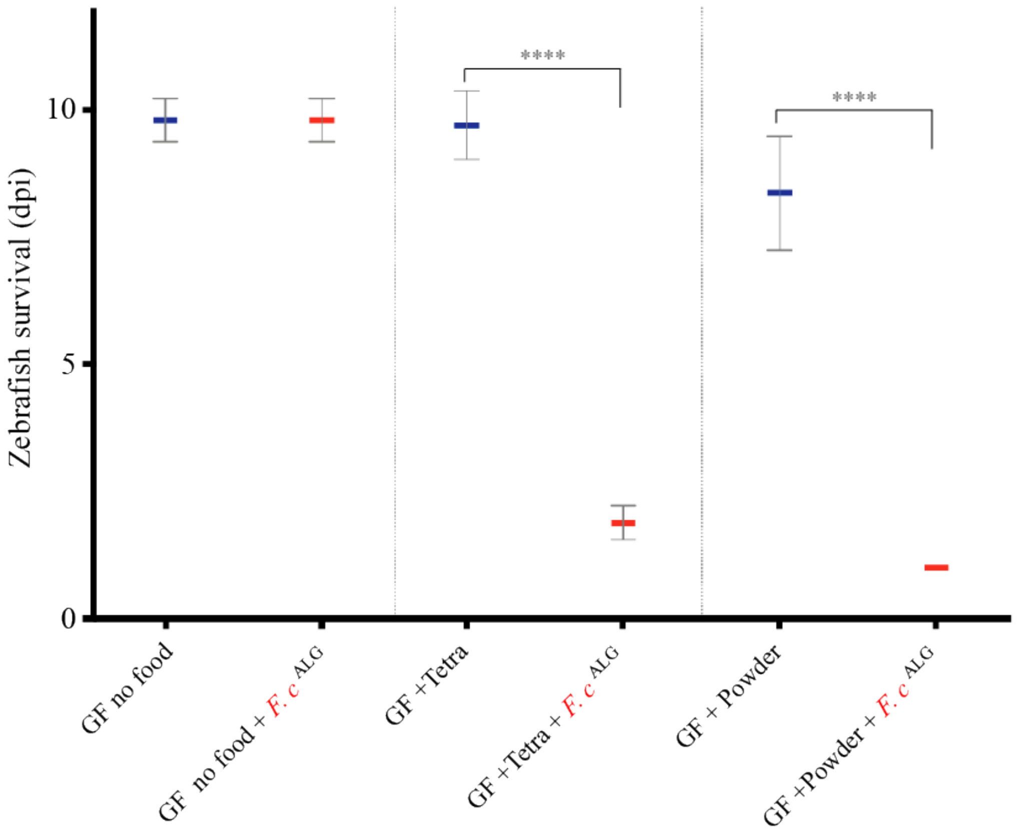


**Supplementary Figure S10. *F. columnare*$^{ALG}$ infection larvae requires food ingestion.** GF zebrafish larvae were fed with sterile *T. thermophila* (GF + Tetra) or sterile fish food powder (GF + powder) or were not fed before *F.columnare*$^{ALG}$ infection. Whereas fed larvae were sensitive to *F.columnare*$^{ALG}$ infection, un-fed GF larvae did not die after fish pathogen infection. Mean survival is represented by a thick horizontal bar with standard deviation. For each condition, n = 12 zebrafish larvae. Blue mean bars correspond to larvae not exposed to the pathogen and red mean bars correspond to exposed larvae. Larvae mortality rate was monitored daily and surviving fish were euthanized at day 10 post infection. Indicated statistics correspond to unpaired, non-parametric Mann-Whitney test. ****: p<0.0001; absence of *: non-significant.

### SUPPLEMENTARY TABLES (Supplementary Table S1-S4).

**Supplementary Table S1. OTUs detected by 16S rRNA clone library generation of bacterial content of 6 and 11dpf old Conv zebrafish larvae, infected or not with *F. columnare*$^{ALG}$.** The number of clones generated are shown for each clone library replicate, a total of 857 clones was generated (minimum clone library coverage 95%).

| | **Conv 6 dpf** | | | **Conv 11 dpf** | | | **Conv 6pf + *F. columnare*** | | | **Conv 11dpf + *F. columnare*** | | |
|---|---|---|---|---|---|---|---|---|---|---|---|---|
| Replicate | R1 | R2 | R3 | R1 | R2 | R3 | R1 | R2 | R3 | R1 | R2 | R3 |
| ***Aeromonas caviae*** | 33 | 3 | 47 | 59 | 31 | 36 | 29 | 4 | 34 | 46 | 75 | 35 |
| ***Aeromonas veronii*** | 8 | 1 | | 4 | 3 | | 3 | 2 | 2 | 11 | 14 | 7 |
| ***Aeromonas veronii 2*** | 16 | 9 | 10 | 7 | 4 | | 11 | 19 | 7 | 11 | 4 | 7 |
| ***Chryseobacterium massiliae*** | | 1 | 2 | 2 | | 2 | 2 | 2 | 2 | 4 | 1 | |
| ***Phyllobacterium myrsinacearum*** | 1 | 2 | 1 | 2 | | 1 | 1 | 1 | 1 | 1 | 2 | |
| ***Pseudomonas sediminis*** | | 5 | 1 | 4 | 1 | | 3 | 2 | 1 | | | 1 |
| ***Pseudomonas mosselii*** | 5 | 4 | 5 | 3 | 6 | 2 | | 12 | 5 | 12 | 3 | |
| ***Pseudomonas nitroreducens*** | 16 | 3 | | 11 | 2 | 12 | 9 | 2 | | 11 | 4 | |
| ***Pseudomonas peli*** | 10 | 19 | 16 | 2 | | | | 2 | 3 | | 2 | 2 |
| ***Stenotrophomonas maltophiliae*** | 1 | 2 | 2 | | 3 | 1 | 2 | | 3 | 1 | | |
| *Delftia tsuruhatensis* | | 2 | | | | | 1 | | | | | |
| *Ensifer adhaerens* | | | | | | | | | | 2 | | |
| *Flavobacterium sp.* | | | | | | | 2 | 1 | 3 | | 1 | |
| *Hydrogenophaga defluvii* | 1 | | | | | | | | | | | |
| *Limnobacter thiooxidans* | | | | 1 | | | | 1 | | | 1 | |
| *Novosphingobium suberraneum* | | 1 | | | | | | | | 1 | | |
| Total number of clones per sample | 91 | 52 | 84 | 95 | 50 | 54 | 63 | 48 | 61 | 100 | 107 | 52 |

**Supplementary Table S2. Percent abundance of OTUs detected by Illumina 16S rRNA sequencing from four other zebrafish facilities for the 10 strains identified as core in the Institut Pasteur zebrafish facility.**
Facility 1 - Hospital Robert Debré, Paris; Facility 2 - Jussieu A2, University Paris 6; Facility 3 - Jussieu - C8 (UMR7622), University Paris 6; Facility 4 – AMAGEN, Gif sur Yvette (commercial facility)

| | **Facility 1** | | **Facility 2** | | **Facility 3** | | **Facility 4** | |
|---|---|---|---|---|---|---|---|---|
| | **6 dpf** | **11 dpf** | **6 dpf** | **11 dpf** | **6 dpf** | **11 dpf** | **6 dpf** | **11 dpf** |
| *Aeromonas caviae* | 0.40 | 0.23 | 0.69 | 8,68 | 0.44 | 0.07 | 33,91 | 0.72 |
| *Aeromonas veronii* | 6,52 | 2,22 | 0.44 | 0.87 | 0.11 | 0.73 | 1,09 | 41,25 |
| *Aeromonas veronii 2* | 0.06 | 0.03 | - | 0.01 | - | - | - | 0.01 |
| *Chryseobacterium massiliae* | 0.51 | 0.08 | 34,83 | 10.79 | 0.003 | - | 28,70 | 2,48 |
| *Phyllobacterium myrsinacearum* | 3,51 | 3,62 | 1,21 | 6,95 | 3,08 | 7,30 | 2,91 | 1,27 |
| *Pseudomonas sediminis* | 0.11 | 0.39 | 1,64 | 1,68 | 0.24 | 0.03 | 0.36 | 0.83 |
| *Pseudomonas mosselii* | 15,30 | 6,51 | 0.02 | 0.11 | 0.38 | 0.66 | 8,12 | 12,19 |
| *Pseudomonas nitroreducens* | 0.17 | 0.95 | 0.40 | 18,41 | 1,59 | 8,87 | 2,66 | 1,75 |
| *Pseudomonas peli* | 39,94 | 53,75 | 29,65 | 13,66 | 15,22 | 38,42 | 2,80 | 2,21 |
| *Stenotrophomonas maltophiliae* | 2,95 | 2,36 | 1,88 | 1,68 | 1,28 | 2,04 | 13,12 | 28,26 |

**Supplementary Table S3.** ***Flavobacterium columnare*** **strains used in this study**

| Strain | Description | Source or Reference | Genomovar |
|---|---|---|---|
| *F. columnare* strains | | | |
| FPC666 | *F. columnare* isolated from *Misgurnus anguillicaudatus* (Japan). | J.F. Bernardet col. | I (Triyanto and Wakabayashi 1999) |
| FPC667 | *F. columnare* isolated from *Carassius auratus* (Japan). | J.F. Bernardet col. | I (Triyanto and Wakabayashi 1999) |
| LD40 07/2489 | *F. columnare* isolated from *Acipenser baeri* (France). | J.F. Bernardet col. | I (this study) |
| LVDI 39/I | UNKNOWN | J.F. Bernardet col. | I (Triyanto and Wakabayashi 1999) |
| JIP 17/01 | *F. columnare* isolated from *Cyprinus carpio* (France). | J.F. Bernardet col. | I (this study) |
| JIP P06/90 | *F. columnare* isolated from *Ictalurus melas* (France). | J.F. Bernardet col. | I (Triyanto and Wakabayashi 1999) |
| LVDL 3414/89 | *F. columnare* isolated from *Anguilla anguilla* (France). | J.F. Bernardet col. | I (Triyanto and Wakabayashi 1999) |
| UJ H2 | *F. columnare* isolated from *Oncorhynchus mykiss* (Finland). | J.F. Bernardet col. | I (this study) |
| UJ B259 | *F. columnare* isolated from outlet water of a rearing tank with infected *O. mykiss* (Finland). | J.F. Bernardet col. | I (this study) |
| UCD Fc4 | *F. columnare* isolated from *O.s mykiss* (USA). | J.F. Bernardet col. | I (Michel, Messiaen, and Bernardet 2002) |
| UCD Fc7 | *F. columnare* isolated from *O. mykiss* (USA). | J.F. Bernardet col. | I (Michel, Messiaen, and Bernardet 2002) |
| JIP P11/91 | *F. columnare* isolated from *O. mykiss* (France). | J.F. Bernardet col. | I (Triyanto and Wakabayashi 1999) |
| JIP 44/87 | *F. columnare* isolated from *Salmo trutta* (France). | J.F. Bernardet col. | I (Michel, Messiaen, and Bernardet 2002) |
| NCIMB2248 | *F. columnare* isolated from *Oncorhynchus tshawytscha* (USA). | J.F. Bernardet col. | I (Triyanto and Wakabayashi 1999) |
| JIP 39/87 | *F. columnare* isolated from *Ictalurus punctatus* (France). | J.F. Bernardet col. | I (Triyanto and Wakabayashi 1999) |
| JIP 07/02(1) | *F. columnare* isolated from *C. carpio* (France). | J.F. Bernardet col. | I/II (this study) |
| ALG-00-530 | *F. columnare* isolated from *Ictalurus punctatus* (USA). | **(Olivares-Fuster et al. 2011)** | II (Arias et al. 2004) |

| | | | |
|---|---|---|---|
| JIP 14/00 | *F. columnare* isolated from *Paracheirodon innesi* (France). | J.F. Bernardet col. | II (Michel, Messiaen, and Bernardet 2002) |
| EK28 | *F. columnare* isolated from *Anguilla japonica* (Japan). | J.F. Bernardet col. | II (Triyanto and Wakabayashi 1999) |
| AJS1 | *F. columnare* isolated from *Poecilia sphenops* (Belgium). | J.F. Bernardet col. | II (Michel, Messiaen, and Bernardet 2002) |
| JIP 13/00 | *F. columnare* isolated from *Paracheirodon innesi* (Hong Kong). | J.F. Bernardet col. | II |
| JIP 02/06(1) | *F. columnare* isolated from *Betta splendens* (Singapore). | J.F. Bernardet col. | II |
| VB1 | *F. columnare* isolated from *Poecilia reticulata* (France). | J.F. Bernardet col. | II (this study) |
| VB2 | *F. columnare* isolated from *Poecilia reticulata* (France). | J.F. Bernardet col. | II (this study) |
| LDA39 H4927 | *F. columnare* isolated from *Ictalurus melas* (France). | J.F. Bernardet col. | III (this study) |
| SNAR C90-106 | *F. columnare* isolated from *Ictalurus punctatus* (USA). | J.F. Bernardet col. | III (Shoemaker and LaFrentz 2015) |
| CIP109753 (PH-97028) | *F. columnare* isolated from *Plecoglossus altivelis* (Japan). | CRBIP | III (Triyanto and Wakabayashi 1999) |
| C#2 | *F. columnare* isolated from *Pelteobagrus fulvidraco* (Unknown). | (Li et al. 2017) | |

Arias, C. R., T. L. Welker, C. A. Shoemaker, J. W. Abernathy, and P. H. Klesius. 2004. "Genetic fingerprinting of Flavobacterium columnare isolates from cultured fish." *J Appl Microbiol* 97 (2):421-8. doi: 10.1111/j.1365-2672.2004.02314.x.

Li, N., Y. Zhu, B. R. LaFrentz, J. P. Evenhuis, D. W. Hunnicutt, R. A. Conrad, P. Barbier, C. W. Gullstrand, J. E. Roets, J. L. Powers, S. S. Kulkarni, D. H. Erbes, J. C. Garcia, P. Nie, and M. J. McBride. 2017. "The Type IX Secretion System Is Required for Virulence of the Fish Pathogen Flavobacterium columnare." *Appl Environ Microbiol* 83 (23). doi: 10.1128/AEM.01769-17.

Michel, C., S. Messiaen, and J.-F. Bernardet. 2002. "Muscle infections in imported neon tetra, Paracheirodon innesi Myers: limited occurrence of microsporidia and predominance of severe forms of columnaris disease caused by an Asian genomovar of Flavobacterium columnare." *Fish Pathology* 25 (5):253-263. doi: https://doi.org/10.1046/j.1365-2761.2002.00364.x.

Olivares-Fuster, O., S. A. Bullard, A. McElwain, M. J. Llosa, and C. R. Arias. 2011. "Adhesion dynamics of Flavobacterium columnare to channel catfish Ictalurus punctatus and zebrafish Danio rerio after immersion challenge." *Dis Aquat Organ* 96 (3):221-7. doi: 10.3354/dao02371.

Shoemaker, C.A. C., and B.R. LaFrentz. 2015. "Lack of association between Flavobacterium columnare genomovar and virulence in hybrid tilapia Oreochromis niloticus (L.) × Oreochromis aureus (Steindachner) " *Fish Pathology* 38 (5):491-498. doi: https://doi.org/10.1111/jfd.12262.

Triyanto, and H. Wakabayashi. 1999. "Genotypic Diversity of Strains of Flavobacterium columnare from Diseased Fishes." *Fish Pathology* 34 (2):65-71. doi: doi.org/10.3147/jsfp.34.65.

**Supplementary Table S4: Combinations of 3, 4, 6 and 7 of the core zebrafish microbiota species tested for their ability to protect against infection by *F. columnare*$^{ALG}$.** These tested combinations did not include *C. massiliae.* None of these tested combinations showed significant protection activity against *F. columnare*$^{ALG}$.

| ***Tested 3 species combinations*** |
|---|
| *A. caviae+A. veronii2+A. veronii1* |
| *P. myrsinacearum+P. mosselli+P. nitroreducens* |
| *P. peli+P. sediminis+S. maltophilia* |
| *A. caviae+A. veronii1+P. myrsinacearum* |
| *A. caviae+A. veronii1+P. nitroreducens* |
| *A. caviae+P. myrsinacearum+P. nitroreducens* |
| *A. caviae+P. nitroreducens+P. sediminis* |
| *A. caviae+P. mosselli+P. peli* |
| *A. caviae+P. peli+S. maltophilia* |
| *A. caviae+P. mosselli+P. nitroreducens* |
| *A. caviae+A. veronii2+P. myrsinacearum* |
| *A. caviae+P. myrsinacearum+S. maltophilia* |
| *A. veronii2+P. mosselli+P. peli* |
| *A. veronii2+P. peli+S. maltophilia* |
| *A. veronii2+P. myrsinacearum+P. nitroreducens* |
| *A. veronii2+P. peli+S. maltophilia* |
| *A. veronii2+P. nitroreducens+P. sediminis* |
| *A. veronii1+P. myrsinacearum+P. mosselli* |
| *A. veronii1+P. mosselli+P. peli* |
| *A. veronii1+P. nitroreducens+S. maltophilia* |
| *A. veronii1+P. myrsinacearum+P. peli* |
| *A. veronii1+P. myrsinacearum+S. maltophilia* |
| *P. myrsinacearum+P. nitroreducens+P. sediminis* |
| *P. myrsinacearum+P. peli+S. maltophilia* |
| *P. mosselli+P. peli+S. maltophilia* |
| *A. caviae+P. nitroreducens+S. maltophilia* |
| *A. veronii2+A. veronii1+P. sediminis* |
| *A. veronii2+P. mosselli+P. peli* |
| *A. caviae+P. peli+P. nitroreducens* |
| ***Tested 4 species combinations*** |
| *A. caviae+A. veronii2+A. veronii1+C. massiliae* |
| *A. caviae+A. veronii2+A. veronii1+P. mosselli* **(non-protecive Mix4 used in Figure 7)** |
| *C. massiliae+P. myrsinacearum+P. mosselli+P. nitroreducens* |
| *C. massiliae+P. peli+P. sediminis+S. maltophilia* |
| *A. caviae+A. veronii2+P. myrsinacearum+P. mosselli* |

| |
|---|
| *A. caviae+A. veronii2+P. nitroreducens+P. peli* |
| *A. caviae+A. veronii2+P. sediminis+S. maltophilia* |
| *P. myrsinacearum+P. mosselli+P. nitroreducens+P. peli* |
| *A. caviae+P. mosselli+P. nitroreducens+P. peli* |
| *A. veronii2+P. mosselli+P. nitroreducens+P. peli* |
| *A. veronii2+P. peli+P. sediminis+S. maltophilia* |
| *P. myrsinacearum+P. mosselli+P. sediminis+S. maltophilia* |
| *A. caviae+A. veronii2+P. mosselli+P. sediminis* |
| *A. caviae+A. veronii2+P. nitroreducens+S. maltophilia* |
| *A. caviae+A. veronii2+P. myrsinacearum+P. peli* |
| *A. caviae+P. myrsinacearum+P. nitroreducens+P. sediminis* |
| *A. veronii2+A. veronii1+P. myrsinacearum+P. mosselli* |
| *A. veronii2+P. myrsinacearum+P. nitroreducens+P. peli* |
| *A. veronii2+P. myrsinacearum+P. sediminis+S. maltophilia* |
| *A. veronii2+P. myrsinacearum+P. peli+S. maltophilia* |
| *A. veronii2+A. veronii1+P. mosselli+P. nitroreducens* |
| *A. veronii2+A. veronii1+P. peli+P. sediminis* |
| *A. veronii2+A. veronii1+P. nitroreducens+S. maltophilia* |
| *A. veronii2+A. veronii1+P. myrsinacearum+P. sediminis* |
| *A. veronii2+A. veronii1+P. myrsinacearum+S. maltophilia* |
| *A. veronii2+A. veronii1+P. mosselli+P. sediminis* |
| *A. veronii2+A. veronii1+P. nitroreducens+P. peli* |
| *A. veronii2+A. veronii1+P. mosselli+S. maltophilia* |
| *A. veronii1+P. myrsinacearum+P. mosselli+P. nitroreducens* |
| *A. veronii1+P. myrsinacearum+P. mosselli+P. peli* |
| *A. veronii1+P. myrsinacearum+P. mosselli+P. sediminis* |
| *A. veronii1+P. myrsinacearum+P. mosselli+S. maltophilia* |
| *A. veronii1+P. nitroreducens+P. peli+S. maltophilia* |
| *A. veronii1+P. peli+P. sediminis+S. maltophilia* |
| *A. veronii1+P. mosselli+P. peli+S. maltophilia* |
| *A. veronii1+P. nitroreducens+P. peli+S. maltophilia* |
| *P. mosselli+A. caviae+P. peli+P. nitroreducens* |
| *A. caviae+P. peli+A. veronii2+P. nitroreducens* |
| ***Tested 6 species combinations*** |
| *A. caviae+P. myrsinacearum+P. nitroreducens+P. peli+P. sediminis+S. maltophilia* |
| *A. caviae+A. veronii1+P. mosselli+P. nitroreducens+P. peli+S. maltophilia* |
| *A. veronii2+A. veronii1+P. mosselli+P. peli+P. sediminis+S. maltophilia* |
| *A. veronii2+A. veronii1+P. myrsinacearum+P. nitroreducens+P. sediminis+S. maltophilia* |
| *A. veronii2+P. myrsinacearum+P. mosselli+P. nitroreducens+P. peli+S. maltophilia* |
| *A. veronii1+P. myrsinacearum+P. mosselli+P. nitroreducens+P. peli+P. sediminis* |
| *A. veronii1+P. myrsinacearum+P. mosselli+P. nitroreducens+P. peli+S. maltophilia* |

| |
|---|
| *A. veronii1+P. mosselli+P. nitroreducens+P. peli+P. sediminis+S. maltophilia* |
| *P. myrsinacearum+P. mosselli+P. nitroreducens+P. peli+P. sediminis+S. maltophilia* |
| *S. maltophilia+A. caviae+P. peli+P. sediminis+P. myrsinacearum+P. nitroreducens* |
| ***Tested 7 species combinations*** |
| *A. caviae+A. veronii2+A. veronii1+P. myrsinacearum+P. mosselli+P. nitroreducens+P. peli* |
| *A. caviae+A. veronii2+A. veronii1+P. myrsinacearum+P. mosselli+P. nitroreducens+P. sediminis* |
| *A. caviae+A. veronii2+A. veronii1+P. myrsinacearum+P. mosselli+P. nitroreducens+S. maltophilia* |
| *A. caviae+A. veronii2+A. veronii1+P. myrsinacearum+P. mosselli+P. peli+P. sediminisa* |
| *A. caviae+A. veronii2+A. veronii1+P. myrsinacearum+P. mosselli+P. sediminis+S. maltophilia* |
| *A. veronii2+A. veronii1+P. myrsinacearum+P. mosselli+P. nitroreducens+P. peli+P. sediminis* |
